# The Art of Not Knowing: Accommodating Structured Missingness in Biomedical Research

**DOI:** 10.1101/2025.06.07.658014

**Authors:** Barbora Rehák Bučková, Charlotte Fraza, Cecilie Koldbæk Lemvigh, Camilla Bärthel Flaaten, Linn Sofie Sæther, Giulia Cattarinussi, Karen Marie Sandø Ambrosen, Ole Andreas Andreassen, Lars Tjelta Westlye, Christian Beckmann, Bjørn Hylsebeck Ebdrup, Paola Dazzan, Torill Ueland, Robin Mitra, Andre Marquand

## Abstract

Missing data remain a ubiquitous and critical challenge in large-scale clinical studies. Despite advances in imputation, most existing methods fail to address structured missingness, where data are missing according a deterministic pattern and which arise due to systematic patterns introduced by experimental design, site protocols, or cohort differences. These patterns violate key assumptions of most imputation algorithms, yet their impact is rarely evaluated.

We demonstrate that structured missingness is a fundamental challenge to drawing valid inferences from standard imputation techniques. First, we present a comprehensive framework for understanding and accommodating the effects of structured missingness. Next, we show through simulations and real-world psychometric data with structured missingness, that widely used algorithms optimised for numerical precision (e.g., Extra Trees, AutoComplete) underperform due to site effects, while donor-based methods (e.g., MICE, hierarchical MICE) better preserve multivariate structure. We propose a novel hierarchical approach that provides optimal performance in simulated and experimental data. Finally, we show that commonly used accuracy metrics, such as mean squared error can obscure these failures, and are therefore inadequate for the evaluation of structured missingness. In contrast, other divergence-based metrics offer a more sensitive and interpretable alternative. We apply this approach to harmonising psychometric data across cohorts, which provides excellent item-level alignment across different instruments.

Our study highlights the need for a paradigm shift in handling missing data within biomedical research, moving beyond conventional imputation frameworks to develop tools that can account for structured missingness. This shift is essential for ensuring reliable inference in multi-site clinical studies, precision medicine, and large-scale population analyses.

## 1 Introduction

Missing data constitute an inherent challenge in biomedical research, arising from a variety of sources, including study design choices, differences in measurement instruments, ascertainment bias, and selective attrition. These factors introduce structured patterns of missingness that, if not taken into account, can bias analyses and limit the generalisability of findings. This issue has become particularly pronounced with the widespread practice of pooling multiple legacy datasets, a strategy that enhances statistical power, improves demographic representation, and enables more sophisticated machine learning applications **[1–3]**. However, cohort aggregation also amplifies structured missingness, as datasets collected under different protocols and for distinct research aims rarely align perfectly in terms of measured variables, participant inclusion, and data completeness. While multi-site studies have become widespread in fields such as psychometrics, multiomics, and neuroscience **[4–6]**, the impact of non-random patterns of missing data in cohort aggregation has not been systematically addressed, particularly in the contexts of data imputation and cohort harmonisation.

A common challenge in the integration of biomedical datasets is that not all variables are collected at every site, leading to systematic patterns of missing data (Fig. 1). An intuitive example of this type of missingness is block missingness, which occurs when data are absent due to study design, site-specific protocols, or measurement differences, rather than at random. For example, in clinical studies, symptoms measures are (by definition) acquired in patients but not controls. Alternatively, in longitudinal studies, follow-up assessments often include different measures than the baseline visit, creating systematic missingness in certain time points or subgroups. This results in deterministic gaps in data tables, where entire sections are systematically missing based on the study site or another aggregation unit. Additionally, block missingness often cooccurs with other missingness patterns, such as dependency missingness (e.g., when missingness of one variable affects missingness of another variable), multimodal linkages (e.g., patients in a study receiving either MRI or PET scans but not both), skip patterns (e.g., survey responses missing because certain answer bypasses follow-up questions), and others **[7]**. Taken together, these patterns fall under the umbrella of structured missingness **[7]** and can severely complicate downstream data analysis.

**Figure 1:**
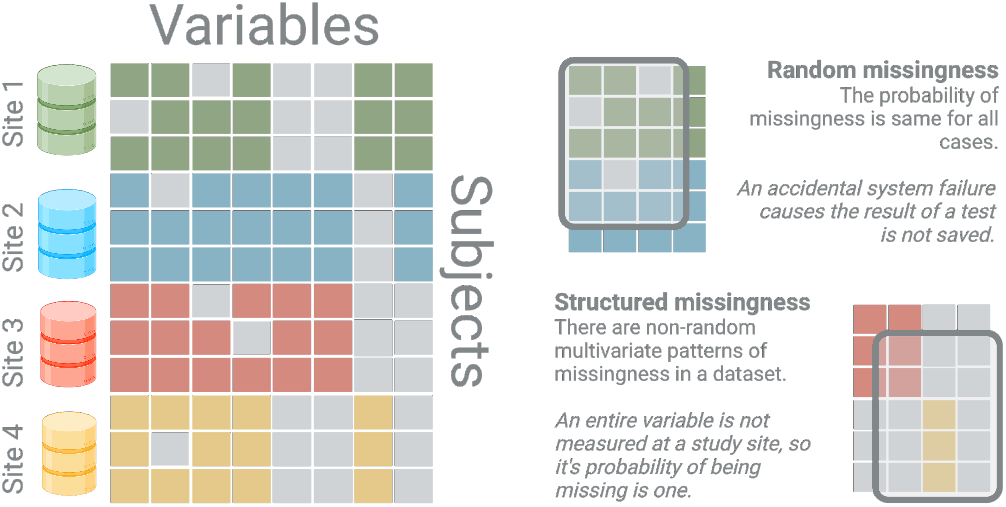
Illustration of structured and random missingness in a dataset. Certain variables are systematically unmeasured at specific study sites, leading to deterministic (probability of missingness equals one) blocks of missing data. This block missingness contrasts with random missingness, which occurs sporadically across the dataset.

Missing data is ubiquitous in biomedical science, yet statistical frameworks for handling missingness have not kept pace with the complexity of today’s integrated datasets. Most imputation methods are still based on Rubin’s 1970s framework, which categorises missing data into three mechanisms: missing completely at random, missing at random, and missing not at random **[8–10]**. Although effective for many single-dataset scenarios **[11–15]**, this framework does not fully account for the systematic gaps introduced when datasets with non-overlapping variables are merged **[16]**.

For instance, missing not at random assumes missingness is influenced by an underlying, unobserved factor, such as patients living farther from a clinic being less likely to return for follow-ups. In this scenario, some patients commuting from a distance still complete their visits, meaning that the missingness is probabilistic rather than absolute. In contrast, block missingness creates deterministic gaps, where entire variables are systematically absent in certain datasets but not others. The limitations of these assumptions become increasingly apparent with current imputation algorithms, which are primarily developed and evaluated for missing completely at random or missing at random scenarios, leaving their effectiveness on data with structured missingness largely unexplored. Many of these algorithms also operate on normality assumptions from which biomedical data often deviate, displaying skewed or multimodal distributions that require more nuanced approaches **[17]**.

Another related, and equally critical issue in multi-site integration involves subtle differences in measurement techniques, which can introduce systematic inconsistencies and obscure biological or clinical insights (Fig. 2) **[18]**. For instance, multi-site clinical trials often measure outcomes like quality of life **[19]** or pain **[20]** using distinct scales, which complicates direct comparisons. Even seemingly well-defined characteristics, such as race **[21, 22]**, socioeconomic status **[23]**, or disease presence **[18]**, can differ across studies, cohorts, and sociodemographic context depending on the specific categories or cut-off values selected. Psychometric data can provide an even more nuanced example: despite the use of standardised constructs, different sites may assess cognition using distinct tests, each with unique scoring methods (i.e., having different distributions) and different sensitivity, specificity, and calibration to underlying latent traits, leading to results that are not comparable between instruments **[24–29]**. These issues can be further intensified by subtle changes in study protocols, recruitment strategies, test administration, language and cultural differences, and data processing pipelines.

**Figure 2:**
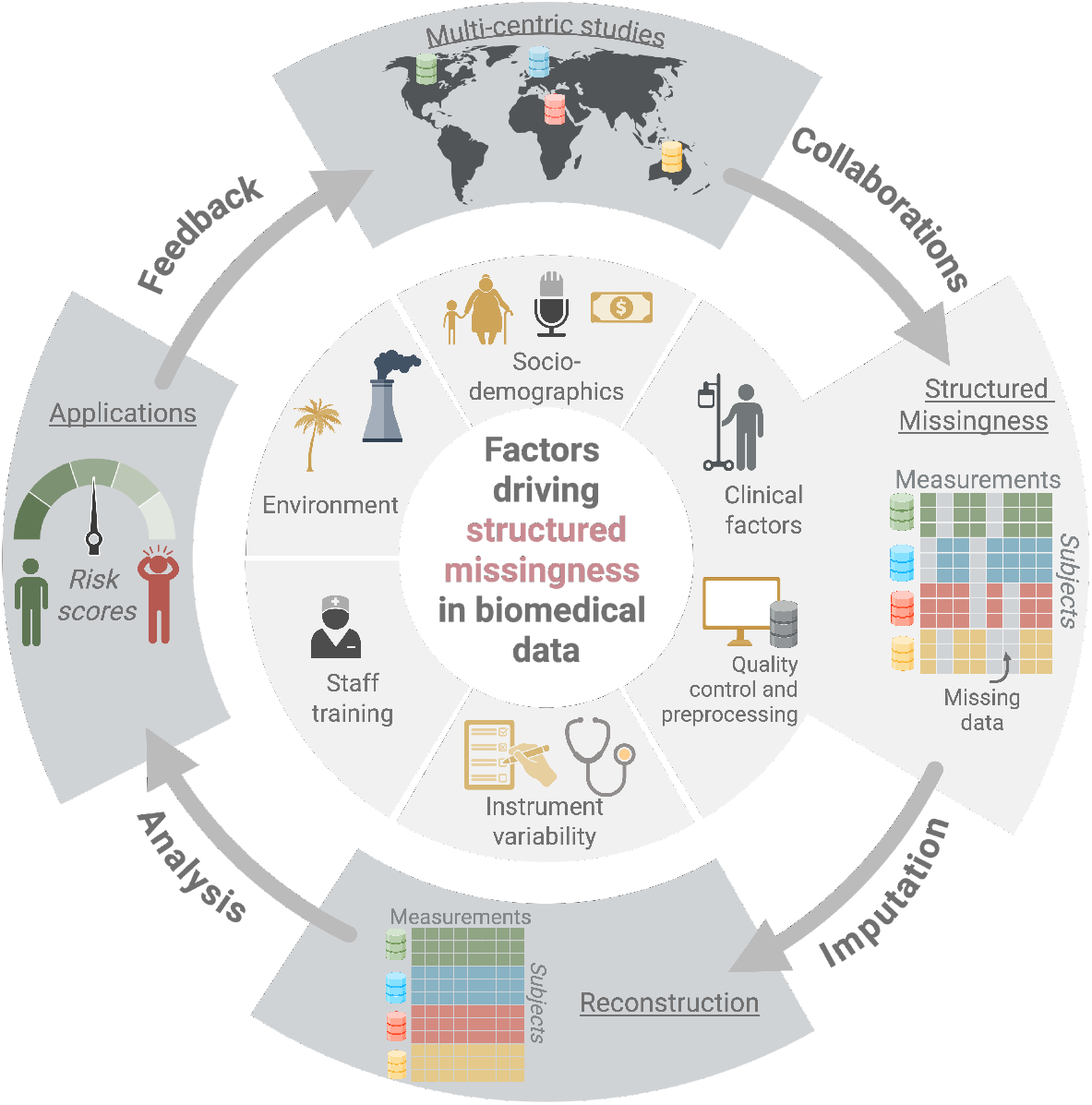
The cycle of collaboration and the factors driving structured missingness in biomedical data-science. Collaborative research often involves merging databases from multiple centres, introducing structured missingness. Missingness may stem from differences in socio-demographic or clinical factors, site-specific data processing, variability introduced by medical devices, test administration, experiment protocols, staff training, or even environmental factors. Once datasets are merged, imputation, harmonisation or variable selection methods are applied to address missingness, ensuring a complete dataset for analysis. The quality of this step is critical, as it directly impacts downstream applications, such as predictive models for identifying patients at risk for certain medical conditions. Finally, the results of the research can further inform acquisition of new measures, support further investigation and strengthen collaborations.

In this work, we argue that addressing structured missingness requires a new framework that is rooted in an extensive simulation of realistic missingness patterns, robust methodologies for imputing data across diverse sources, and rigorous evaluation to ensure that integrated analyses yield reliable and interpretable insights. To achieve this, we: (i) Conduct a large-scale simulation study to systematically assess the impact of structured missingness on the performance of state-of-the-art imputation algorithms. (ii) Demonstrate that widely used imputation methods fail systematically under structured missingness and propose a comprehensive framework for evaluating imputation approaches. We introduce evaluation metrics that capture the complexities of structured missingness, moving beyond traditional single-dataset benchmarks to a multi-faceted assessment based on diverse statistical properties. (iii) Apply our framework to a large multi-site study on cognition in psychosis, focusing on the challenges of imputing structured missing data across multiple sites with different protocols and data collection practices. (iv) Discuss key considerations for real-world applications, including the integration of heterogeneous datasets with non-overlapping variables, and demonstrate how our approach improves imputation performance in a custom dataset of cognitive assessments.

## 2 Results

### 2.1 Simulation

To systematically evaluate the impact of structured missingness on imputation, we designed an extensive simulation framework that reflects the complexity of real-world data (Fig. 3). Having access to the ground truth (the original complete data before any values are removed) enables us to quantify the effect of structured missingness on imputation accuracy, identify the most informative evaluation criteria, and ultimately determine the most suitable imputation approach.

**Figure 3:**
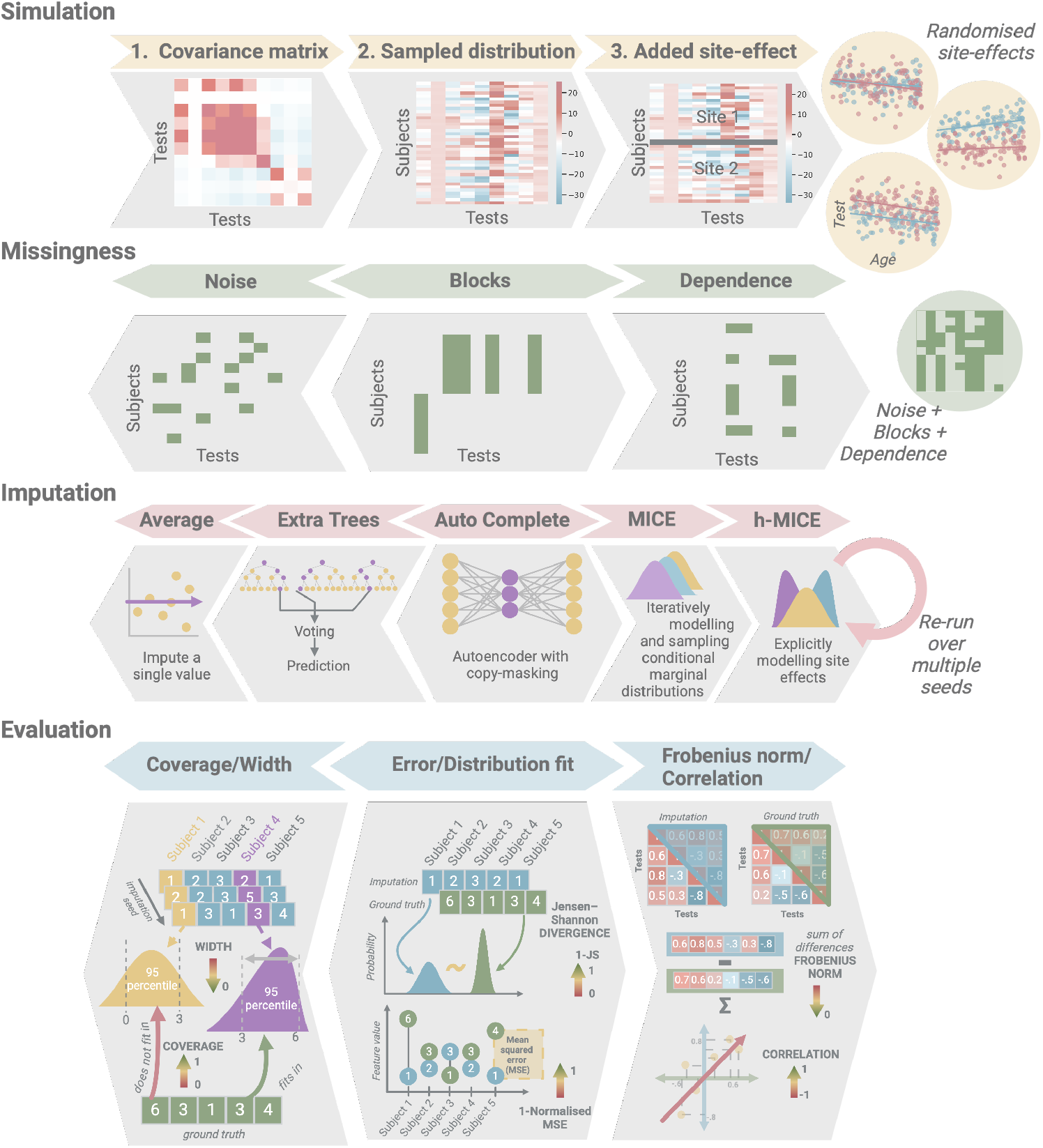
Simulation study pipeline: The first row illustrates the data generation process, starting with the extraction of a covariance matrix from a real-world dataset, generating synthetic datasets, and introducing site effects through random linear offsets (yellow circles). The second row depicts the three types of missingness introduced: missing completely at random, block missingness (entire variables deleted for a site), and dependence missingness (missingness in one variable driven by another). The missingness patterns were generated individually and in combination (green circle). The third row shows the imputation algorithms applied to the incomplete datasets: Average imputation, Extra Trees, AutoComplete, the vanilla MICE, and hierarchical MICE, each of which was sampled ten times. The final row shows the quality of imputation using subject-level metrics (coverage and width), variable-level metrics (1 − negative mean squared error and 1− Jensen-Shannon divergence), and multivariable relationship metrics (Frobenius norm and upper-triangle correlation of correlation matrices).

First, we extracted the covariance matrix from a real-world dataset of cognitive measures (see Data description in the Methods section, or the GitHub repository) to ensure realistic multivariate relationships. Using this as the basis for a multivariate normal distribution, we generated two synthetic datasets, each containing 40 variables, with sample sizes of either 100 or 1,000 subjects. To simulate site effects, we introduced random linear offsets across two predefined sites, with the magnitude of these offsets varying randomly across datasets and variables (for further details, refer to the Methods section).

Next, we modelled three types of missingness: (i) *Missing completely at random with no structure*, by randomly removing values from randomly selected variables to ensure the absence of structured patterns; (ii) *Block missingness*, where entire variables were systematically absent at one of the two sites, an instance of strong structured missingness; and (iii) *Dependence missingness*, in which the presence of missing values in one variable was conditionally dependent on another. While this list is not exhaustive (see **[16]** for a more comprehensive typology), it reflects several common scenarios encountered when merging heterogeneous datasets. In the case of block missingness, a key factor influencing imputation quality is the degree of overlap between datasets. To assess this, we varied the number of shared variables between sites, testing overlaps of 4, 10, and 20 variables. Given the complexity of real-world missingness patterns, we simulated these mechanisms both individually and in combination.

To recover incomplete datasets, we applied four state-of-the-art imputation algorithms (Table 1), each representing a distinct computational approach to handling missing data. These included the Extra Trees algorithm, which has shown promise in real-world applications by balancing numerical accuracy with the preservation of the shape of the underlying distribution **[6]**. We implemented the recently published AutoComplete method, a state-of-the-art autoencoder that has demonstrated strong performance on genetic datasets **[30]**. Furthermore, we included the widely used Multiple Imputation by Chained Equations (MICE) **[31]**, a standard approach for handling missing data. Additionally, we developed a hierarchical extension of MICE that explicitly models site-specific offsets (systematic differences in variable distributions arising from multi-site data collection) **[32]**. By estimating offsets only when a variable is observed at multiple sites, this method preserves inter-site variability without overfitting to single-site data. This hybrid strategy enables improved coherence across the integrated dataset while respecting the local structure of site-specific measurements. Finally, we benchmarked all methods against simple average imputation. Although we do not recommend this approach, it is widely used in practice for its simplicity and computational efficiency, so we include it to illustrate its shortcomings and highlight the need for more advanced methods.

**Table 1:**
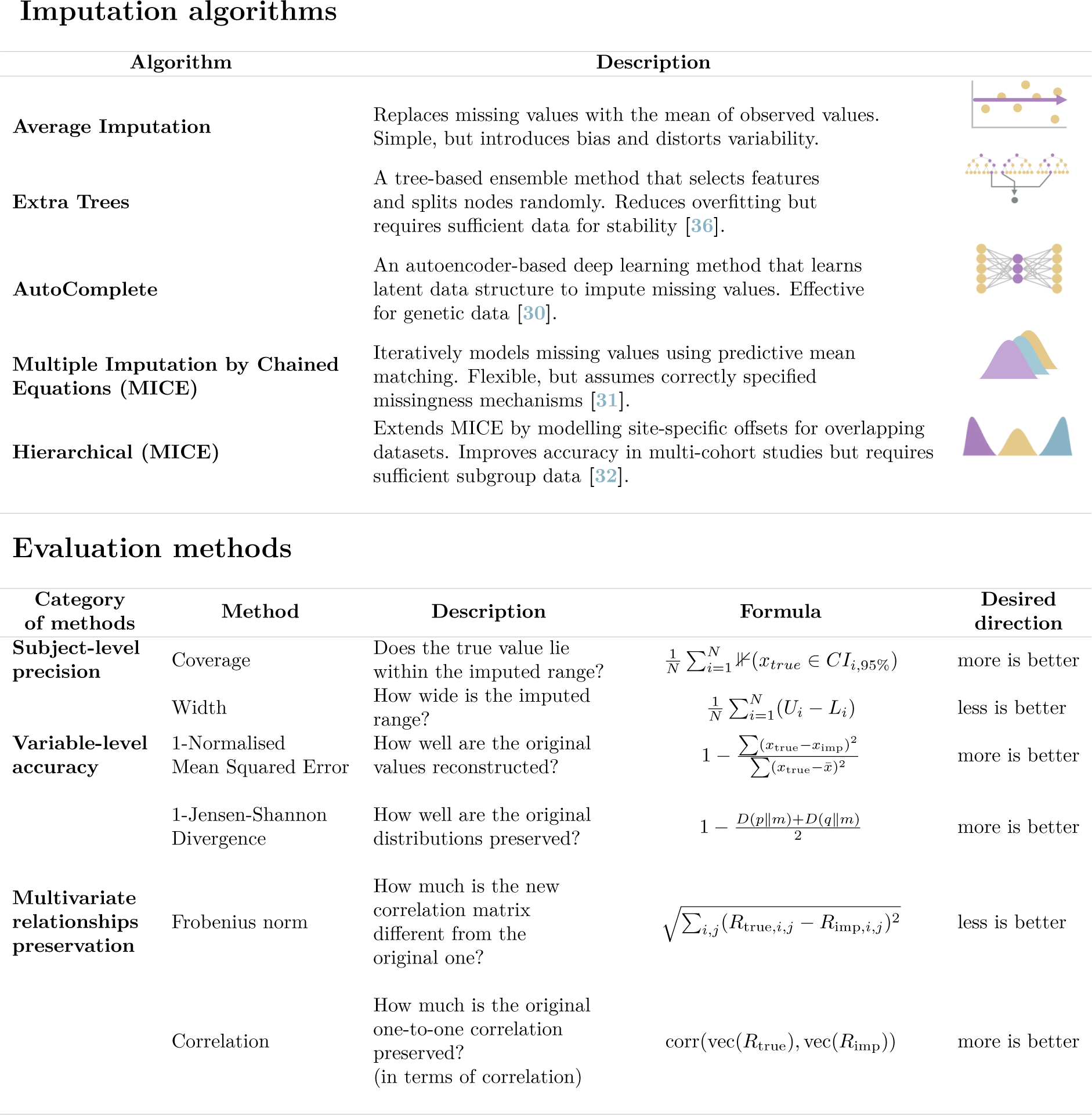
Overview of imputation algorithms and evaluation metrics used in this study. The table summarizes the imputation algorithms, ranging from simple mean substitution to advanced machine learning and hierarchical methods, along with their descriptions. The evaluation metrics assess imputation quality in three domains: subject-level precision, variable-level accuracy, and preservation of multivariate relationships. For each metric, we provide a description, formula, and the desired performance direction. *Parameter descriptions:* In Coverage, *x*_*t*_*rue* is the true value, *CI*_*i*,95%_ is the 95% confidence interval, and *N* is the total number of imputation seeds. In Width, *U*_*i*_ and *L*_*i*_ are the upper and lower bounds, respectively. In 1-Normalised MSE, *x*_true_ and *x*_imp_ are the true and imputed values, and 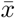 is the mean of the true values. In 1-Jensen-Shannon Divergence, *p* and *q* are the original and imputed distributions, and *D*(· ∥ ·) is the Kullback-Leibler divergence. In Frobenius norm, *R*_true_ and *R*_imp_ are the true and imputed correlation matrices, and|| *·* || _*F*_ is the Frobenius norm. In Correlation, corr(·, ·) is the correlation between the vectorized matrices.

Imputation algorithms
| Algorithm | Description |
| --- | --- |
| Average Imputation | Replaces missing values with the mean of observed values. Simple, but introduces bias and distorts variability. |
| Extra Trees | A tree-based ensemble method that selects features and splits nodes randomly. Reduces overfitting but requires sufficient data for stability [36]. |
| AutoComplete | An autoencoder-based deep learning method that learns latent data structure to impute missing values. Effective for genetic data [30]. |
| Multiple Imputation by Chained Equations (MICE) | Iteratively models missing values using predictive mean matching. Flexible, but assumes correctly specified missingness mechanisms [31]. |
| Hierarchical (MICE) | Extends MICE by modelling site-specific offsets for overlapping datasets. Improves accuracy in multi-cohort studies but requires sufficient subgroup data [32]. |

Table 1: Overview of imputation algorithms and evaluation metrics used in this study.
| Category of methods | Method | Description | Formula | Desired direction |
| --- | --- | --- | --- | --- |
| Subject-level precision | Coverage | Does the true value lie within the imputed range? | $\frac{1}{N} \sum_{i=1}^N \mathbb{I}(x_{true} \in CI_{i,95\%})$ | more is better |
| | Width | How wide is the imputed range? | $\frac{1}{N} \sum_{i=1}^N (U_i - L_i)$ | less is better |
| Variable-level accuracy | 1-Normalised Mean Squared Error | How well are the original values reconstructed? | $1 - \frac{\sum (x_{true} - x_{imp})^2}{\sum (x_{true} - \bar{x})^2}$ | more is better |
| | 1-Jensen-Shannon Divergence | How well are the original distributions preserved? | $1 - \frac{D(p\vert \vert m) + D(q\vert \vert m)}{2}$ | more is better |
| Multivariate relationships preservation | Frobenius norm | How much is the new correlation matrix different from the original one? | $\sqrt{\sum_{i,j} (R_{true,i,j} - R_{imp,i,j})^2}$ | less is better |
| | Correlation | How much is the original one-to-one correlation preserved? (in terms of correlation) | $\text{corr}(\text{vec}(R_{true}), \text{vec}(R_{imp}))$ | more is better |

A notable difference among the tested algorithms is their ability to quantify their own uncertainty - essentially, how confident they are in the values they impute. Methods like MICE inherently capture uncertainty by sampling from a fitted posterior distribution, whereas tree- or neural-network-based approaches, such as Extra Trees and AutoComplete, do not naturally offer this. Given that the stability of imputation is crucial for reliable evaluation, we performed each imputation ten times using different random seeds. This allowed us to assess variability in the imputed values and ensure consistency across repeated runs.

Imputation procedure is often optimised and assessed using a single metric, such as mean squared error that focuses solely on pointwise reconstruction accuracy **[33]**. However, this approach can be misleading; for instance, mean imputation may minimize average error while distorting the underlying data distribution, thereby compromising downstream analyses **[34, 35]**. This problem can be exacerbated in the presence of structured missingness, where blocks of missing data and site-specific effects make it nearly impossible to accurately estimate the magnitude and direction of bias. Consequently, methods focused solely on numerical accuracy may offer a biased and incomplete picture of the performance. To provide a more comprehensive assessment and identify criteria more robust towards structured missingness, we adopted a multi-dimensional evaluation framework.

We evaluate these methods at three levels, namely at the level of subjects, individual variables, and multivariate relationships (Fig. 3). At the subject level, we measured *coverage*—whether the true value falls within the imputed range, and *interval width* to guard against excessively broad uncertainty estimates. At the variable level, we computed the *normalised mean squared error* (1̆*NMSE*) to assess precision and *Jensen–Shannon divergence* (1̆*JSD*) to assess the preservation of the original data distribution. Finally, because many downstream analyses rely on multivariate relationships rather than individual values, we assessed the preservation of these relationships using the *Frobenius norm* and the *correlation* of the upper triangle of the correlation matrix of the reconstructed data, capturing both, overall correlation structure and potential shifts in pairwise relationships.

Applying these metrics to our simulated datasets revealed substantial differences in performance across imputation methods (Fig.4). Even under random missingness, the performance of most methods was suboptimal, and significantly deteriorated with the introduction of structured missingness (Fig.4). In the random missingness condition (first column of Fig.4), MICE-based algorithms outperformed other methods at the subject level, achieving coverage close to 0.8, although at the cost of wider uncertainty intervals compared to Extra Trees and AutoComplete. At the variable level, all methods outperformed simple mean imputation in terms of normalised mean squared error, though MICE-based methods showed notably high variability in quality, which decreased with larger sample sizes. In terms of distribution preservation (quantified by 1 − JSD), all algorithms performed comparably, except for standard Auto-Complete, which struggled with larger sample sizes, and mean imputation, which collapsed the original data distribution to a single value (i.e., a spike in the distribution). At the multivariate level, hierarchical MICE best preserved relationships across variables, outperforming other methods in both Frobenius norm and correlation-based metrics. Overall, under the random missingness assumption (for which these algorithms were originally designed), the most notable performance differences appeared at the subject level, with MICE-based methods, particularly hierarchical MICE, demonstrating the most consistent results across all evaluation levels.

**Figure 4:**
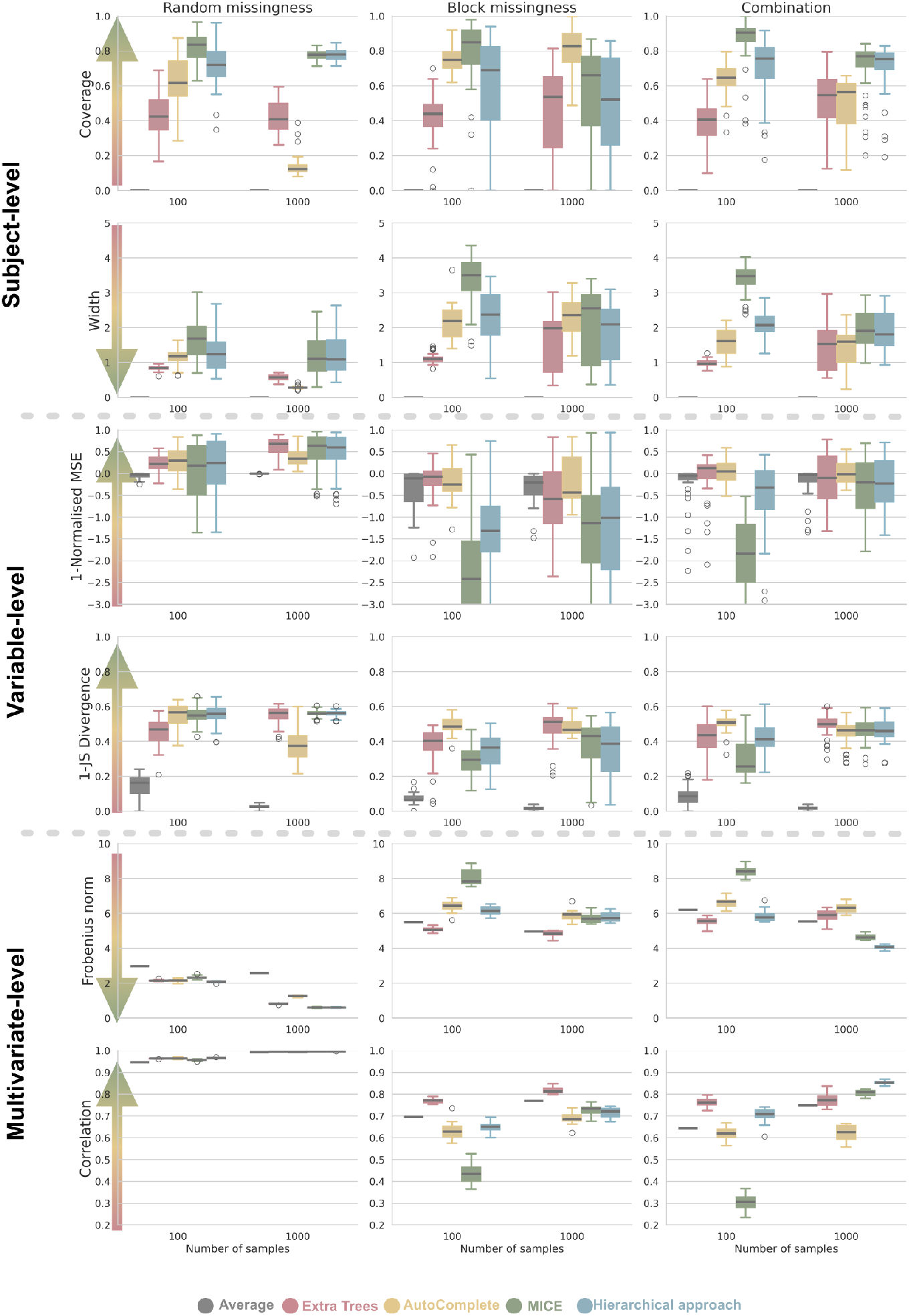
Evaluation of the imputation methods on the simulated datasets: Each row represents a quality of imputation criterion, each column represents a different type of missingness. The results are shown for five imputation algorithms: Average imputation, Extra Trees, AutoComplete, MICE, and hierarchical MICE; and for two different dataset sizes (100, 1000). Here, the amount of random noise missingness was set to 30%, while both batteries overlapped in 10 variables. The arrows on the left side point in the direction of the ideal reconstruction values.

Introducing blockwise missingness (second column of Fig. 4) led to a marked increase in uncertainty across all methods, as evident by the widened boxplots and whisker ranges. At the subject level, AutoComplete outperformed standard MICE at higher sample sizes, while hierarchical MICE showed a notable decline in performance. Extra Trees maintained its average performance but exhibited greater variability in imputed values. At the variable level, both standard and hierarchical MICE struggled with NMSE, performing significantly worse than in the random missingness scenario. However, Extra Trees and AutoComplete also performed poorly, even worse than mean imputation. Multivariate relationships were also affected, as indicated by increased Frobenius norm values. Standard MICE, in particular, failed to maintain above-average performance in smaller samples, whereas hierarchical MICE and Extra Trees better preserved multivariate structures. Finally, when blockwise and random missingness were combined, results fell between the two extremes, with overall performance largely determined by the relative proportions of each missingness type.

The additional scenarios presented in Appendix show trends consistent with those described above. Regardless of the size of the overlap or the percentage of random missingness, the overall patterns remain stable. In general, a lower degree of missingness leads to better imputation performance when combined with block missingness. Furthermore, the size of variable overlap between datasets plays a crucial role in preserving multivariate relationships: Smaller overlaps result in greater deterioration of these relationships.

### 2.2 Real-world data analysis

As a proof-of-concept application for this framework for missing data, we applied it to two problems of increasing complexity. First, we imputed data of healthy controls from two overlapping cognitive batteries from the Norwegian ‘Thematically Organised Psychosis’ (TOP) cohort **[37]**. The dataset comprised two cognitive test batteries of 509 and 667 subjects, with 22 overlapping scores and a set of additional unique scores for each battery (11 and 5, respectively). In this case, the structured missingness arose because the cognitive battery was revised and substituted with another part-way through the acquisition. This was done to reduce the burden on participants in that the second battery was designed to measure similar constructs more time-efficiently. This is an ideal configuration to validate our approach, as other factors that might introduce variability between datasets (e.g. demographic or cohort differences) were effectively controlled for by the study design.

In the second scenario, we incorporated three additional datasets collected at the Centre for Neuropsychiatric Schizophrenia Research (CNSR) in Denmark **[38–40]**. These datasets were acquired sequentially as part of various studies (comprising 80, 119 and 100 subjects), capturing mostly, but not entirely, overlapping variables. Introducing additional datasets further challenged the imputation algorithms. First, site effects were expected to be more pronounced, as data collection occurred in a different language (implying a translated variant of the test), with different staff, and on a distinct cohort. At the same time, integrating additional datasets increased overall diversity, which could ultimately improve downstream analyses, as statistical and machine learning algorithms fundamentally rely on identifying and leveraging multivariate patterns in the data.

To evaluate imputation performance against ground truth, we first masked the original merged dataset 20 times. Specifically, in each fold, we randomly masked out 5% of each variable’s values (without repetition), on top of the values that were already missing. This ensured that, alongside imputing originally missing values, we also imputed a subset of values for which we had ground-truth data. After all iterations, we could fully reconstruct the original dataset, enabling a direct comparison between imputed and true values. As in the simulation study, each algorithm was initialised ten times per mask to assess variability in imputation outcomes. Finally, we computed evaluation metrics based on the ground-truth values (Fig. 5).

**Figure 5:**
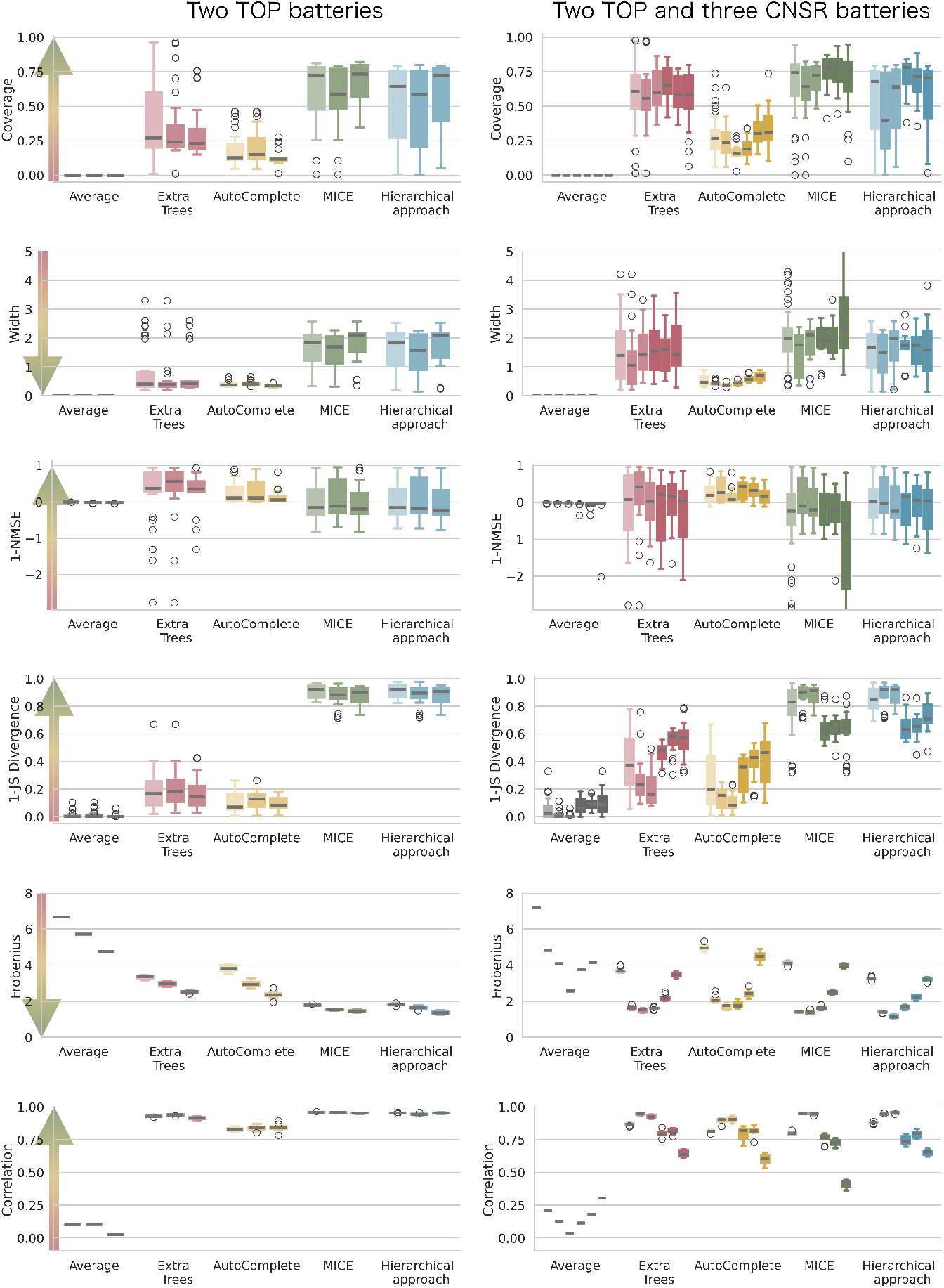
Evaluation of k-fold cross-validation on a real-world dataset: Each subplot presents box-plots for different imputation algorithms, color-coded for distinction. Within each algorithm, the first (lighter-colored) boxplot represents the quality measure computed on the entire dataset, while subsequent boxplots correspond to individual sites. The left panel shows results for the analysis using only the two TOP cognitive batteries, while the right panel extends the analysis to include three additional CNSR datasets (shown in darker colours). Arrows on the left indicate the direction of ideal reconstruction values.

Overall, the imputation results aligned closely with our simulation findings (left column of Fig. 5). Subject-specific metrics, such as coverage and interval width, were dominated by MICE-based approaches, whereas Extra-Trees and Auto-Complete produced imputed values with a much narrower range, failing to capture the true values. Consequently, these methods outperformed MICE-based approaches in the normalised mean squared error metric. The most notable deviation from our simulations was the extent to which MICE-based methods preserved the original data distribution. This advantage was further reflected in the preservation of multivariate relationships, particularly as measured by the Frobenius norm. Furthermore, as data from additional sites were integrated (right column of Fig. 5), imputation quality for the original two batteries did not significantly change. This resilience is encouraging, as it demonstrates that the imputation for the original datasets was not negatively perturbed by the new sources of variance introduced by the additional data. However, these evaluations only captured how well the algorithms reconstructed values that were originally observed without providing insight into the plausibility of imputations for values that had been missing from the outset.

To evaluate imputation plausibility, we used JSD to compare the distribution of imputed values to that of the observed data for each variable. Ideally, in the absence of structured missingness, the imputed distribution should closely resemble the observed data. In cases where a variable was measured at both sites but exhibited site effects, an optimal imputation method would ensure that the imputed density at each site aligns with the respective observed density. However, as shown in the left column of Fig. 6B, some methods struggled with this task, particularly AutoComplete, which produced overly narrow priors that were incompatible with the original distribution. For variables measured at only one site, an effective imputation method should (i) preserve the original distribution at the observed site and (ii) generate a plausible distribution at the unobserved site, informed by the available data from other locations. The goal is not to produce an identical match but rather a reasonable approximation that accounts for site-specific variation. Fig. 6B illustrates the varying degrees of success across different algorithms in addressing this challenge.

**Figure 6:**
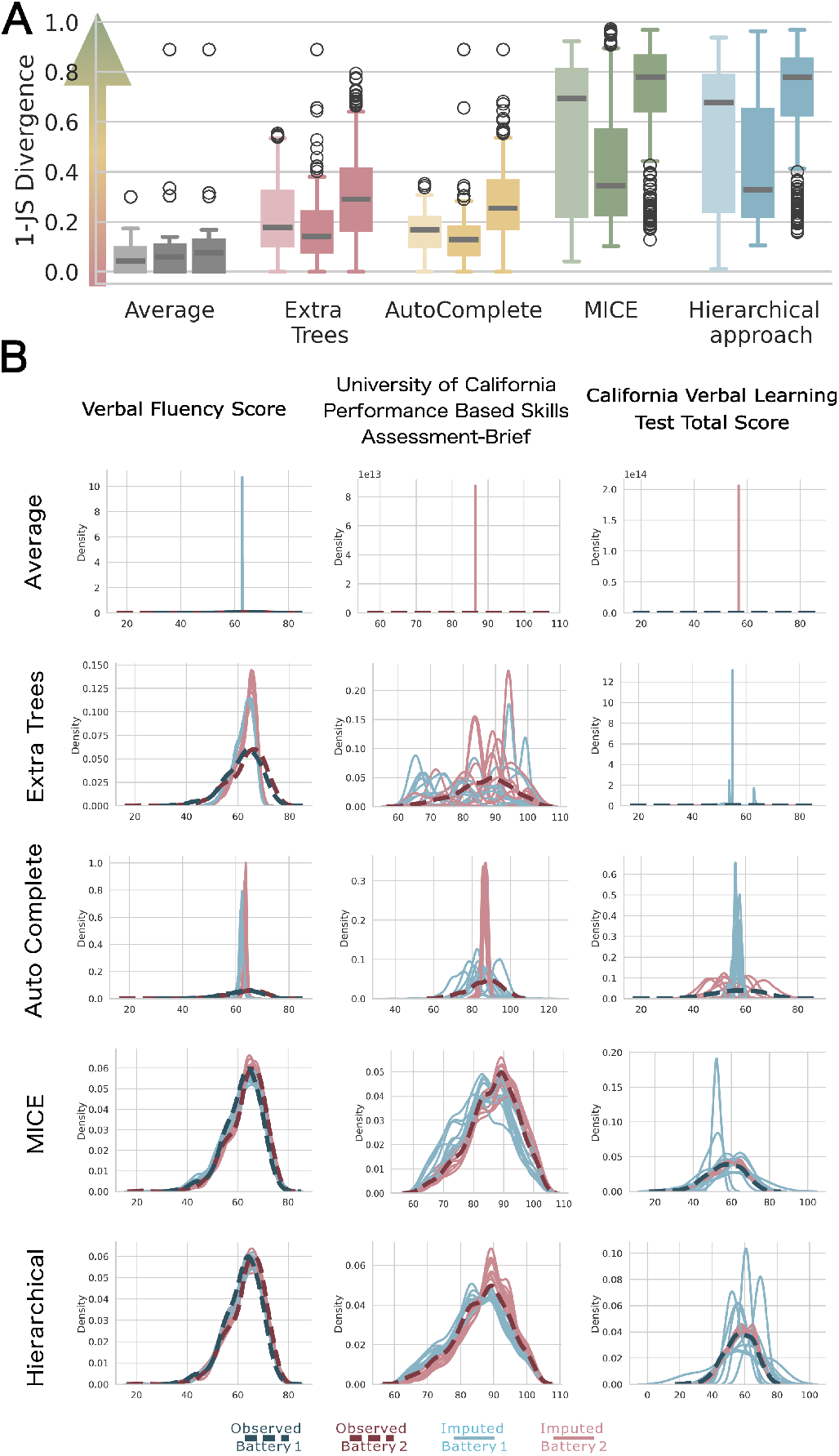
(A) Comparison of the imputed variable distributions with the original distributions, quantified by 1− Jensen Shannon divergence. (B) Examples of imputation approaches (rows) for cognitive variables (columns) across the two batteries of the TOP dataset—dark dashed lines represent the distribution of the original data, while thin lines indicate the distribution of the imputed data. In the first column, the variable was a part of both TOP batteries, while in the second and third column, the variable was only measured at red and blue site, respectively.

Both the standard MICE and the Hierarchical method outperformed the alternative approaches (Fig. 6). Panel B of Fig. 6 clearly illustrates the shortcomings of individual approaches. The first column of density plots shows that most methods reconstructed the variable well when measured at both sites. However, in the other columns, where only one site measured the variable, Extra-Trees performed poorly, with highly variable distributions fitted at the unmeasured site. Auto-Complete, in contrast, adopted a narrow normal distribution to optimise absolute error, while MICE and the hierarchical method performed well, reflecting the distribution of the measured site. These figures show that despite an apparent equivalence in the summary statistics, the imputations produced by the hierarchical MICE are clearly superior to the standard implementation, showing much smoother distributions that have a better correspondence to the true distribution for the unmeasured site.

## 3 Discussion

We present a comprehensive framework for imputation and assessment of structured missingness and show that: (i) Evaluation across multiple metrics is essential to obtain an accurate assessment of the performance of different imputation algorithms; (ii) Even algorithms that are regarded to perform well in general can fail in the context of structured missingness; (iii) Overall, MICE based methods provided the best balance between coverage and accuracy metrics and (iv) accommodating hierarchical structure in the imputation can improve performance, particularly for preserving multivariate relationships.

Our results indicate that some approaches, like donor-based matching in MICE, are by design better suited to structured missingness than others, such as deep learning models or decision forests. While Extra-Trees and Auto-Complete performed well in previous benchmarks **[6, 30]**, their reliance on numerical precision as the primary objective limited their effectiveness in the presence of structured missingness. Extra-Trees, for instance, produced highly variable imputations when site effects were present, failing to capture the underlying data structure (Fig. 6). Auto-Complete, by contrast, defaulted to narrow normal distributions, leading to overly confident predictions that did not reflect true variability. However, although these models are not ideally suited in their current form, future adaptations could improve their capabilities, for instance, by prioritising Kullback–Leibler divergence as an optimisation criterion in autoencoders, or site-aware regularisation in Extra Trees. However, progress in improving imputation methods will require moving beyond conventional accuracy metrics like mean squared error and considering evaluation strategies that capture structured missingness effects.

Our findings further highlight the critical role of preserving distributional properties and multivariate relationships when using imputation methods under structured missingness. Indeed, methods that maintained these properties, particularly our hierarchical MICE approach, demonstrated the preservation of both, thereby improving the robustness of the potential downstream analyses. As shown in Fig. 6, the hierarchical MICE method consistently produced imputations that better reflected the distribution of the measured site, even when entire variables were missing from a given location. These results underscore the importance of imputation strategies that do not optimise solely for a numerical error measure such as mean squared error. Our analysis of data from real-world cognitive batteries further underscores both the potential and limitations of imputation in addressing structured missingness. When site effects were minimal due to consistent data collection protocols, we were able to isolate the significant impact of structured missingness caused by the replacement of test batteries. In this setting, our hierarchical approach demonstrated its strength in preserving distributional and multivariate properties. As we expanded the analysis by incorporating additional datasets with more pronounced site effects, we observed a slight adjustment in performance for the initial datasets (Fig. 5). However, overall performance remained stable or even improved for most imputation methods, including the best-performing hierarchical approach.

Furthermore, our results highlight that conventional evaluation metrics, such as numerical precision, can be misleading under structured missingness. While all metrics reflected the impact of structured missingness, those focused on numerical precision, such as coverage, interval width, and normalised mean squared error, were disproportionately sensitive to site effects. In particular, when systematic shifts in means occurred across datasets, these methods failed to capture the true underlying distribution. This resulted in artificially low error measures even when the imputed values did not match the original structure. For example, if a site effect causes an offset in a variable, an imputation algorithm might correctly reconstruct individual values within one site while shifting them toward the mean of another. In such cases, 1 −NMSE would be negative and coverage would be low, even though the algorithm successfully preserved the general distribution of the variable. This example underscores a fundamental limitation of conventional error metrics: when structured missingness is present, optimising for numerical accuracy alone can degrade the integrity of the dataset. Indeed, as seen in our simulations, mean imputation will *by definition* achieve a lower normalised mean-square error while severely distorting the original distribution (Fig. 4).

The issue of misleading evaluation metrics becomes more pertinent with the increasing prevalence of multi-centre studies and data-sharing initiatives. As structured missingness arises across biomedical research due to site-specific assessment protocols, it is likely more widespread than previously recognised, and if unaddressed, may lead to systematic biases in multi-site datasets. This risk is particularly pronounced in precision medicine, where biased imputations can influence treatment recommendations. For example, in pharmacogenomics, systematic errors in imputing missing genotype data can lead to incorrect dosage recommendations, potentially compromising patient safety **[41, 42]**. Furthermore, as demonstrated in our real-world example, it hinders the use of cognitive data across populations, and, when combined with neuroimaging measures, can obscure relationships between brain structure (or function) and behavioural outcomes. Multi-omics studies also face these challenges, as differences in platforms or batch effects systematically introduce missing data within or across sites. Consequently, these findings should be considered a starting point for further research into structured missingness and appropriate methods for mitigating its effects across various disciplines.

Although our approach to simulating block missingness effectively captures a subset of real-world missingness patterns, it does not encompass the full complexity of structured missingness encountered in practice, such as batch effects, skip patterns, multimodal linkage and others outlined in Mitra et al **[7, 16]**. A deeper understanding of structured missingness will enable better integration of latent factors, variable dependencies, and dynamic missingness mechanisms, ultimately improving imputation strategies, as well as evaluation criteria **[43, 44]**. However, even with improved theoretical understanding, bridging the gap between theory and the practical implementation of imputation methods across heterogeneous datasets remains a challenge **[45]**.If missingness patterns reflect demographic, socioeconomic, or systemic factors, employing imputation methods that ignore these disparities risks reinforcing biases, skewing analyses, and ultimately exacerbating health inequities **[46]**. Moreover, if datasets differ in their demographic makeup, where one dataset predominantly includes older adults while another focusses on adolescents, imputation models may not be able to generalise effectively, resulting in biased predictions and potentially flawed inferences **[47]**. Addressing these challenges requires thoughtful preprocessing and model design to mitigate risks of bias.

While imputation is the initial step of the data-analytical pipeline that recovers missing values in a form that preserves site-specific variability, meaningful analysis across multiple datasets requires additional steps to account for systematic site effects. Harmonisation enables meaningful comparisons across sites by transforming data into a common space. In many multi-site studies, adjusting for site effects is sufficient for group-level comparisons **[4–6]**. However, these transformations inevitably introduce uncertainty and reshape the dataset in ways that traditional modelling approaches may not fully accommodate. Imputation itself generates multiple plausible values per missing data point, whether it is in probabilistic approaches, where posterior distributions captures inherent uncertainty, or more deterministic approaches, where the algorithmic uncertainty can be estimated through multiple random initialisations. However, most machine learning algorithms are not designed to handle multiple versions of the imputed dataset, limiting the reliability of downstream analyses. Addressing this challenge requires methods that explicitly account for uncertainty and site variability.

Normative modelling **[48, 49]** is particularly well-suited for this purpose. Rather than focusing on reconstructing absolute values, normative modelling emphasises preserving the underlying data distributions and multivariate relationships, making it robust to the uncertainties introduced by imputation and harmonisation. By constructing a universal population-wide model that can be adjusted for individual sites, normative modelling leverages the increased sample size while accounting for site-specific effects (Fig. 7). This approach is particularly advantageous when precise individual imputations are unreliable; instead, group-level imputations can recover the overall data structure, and predictions at the individual level are made only when data are actually observed. In doing so, normative modelling not only preserves biologically meaningful variability and risk indicators **[1, 50]**, but also avoids the pitfalls of forcing uniformity across heterogeneous datasets.

**Figure 7:**
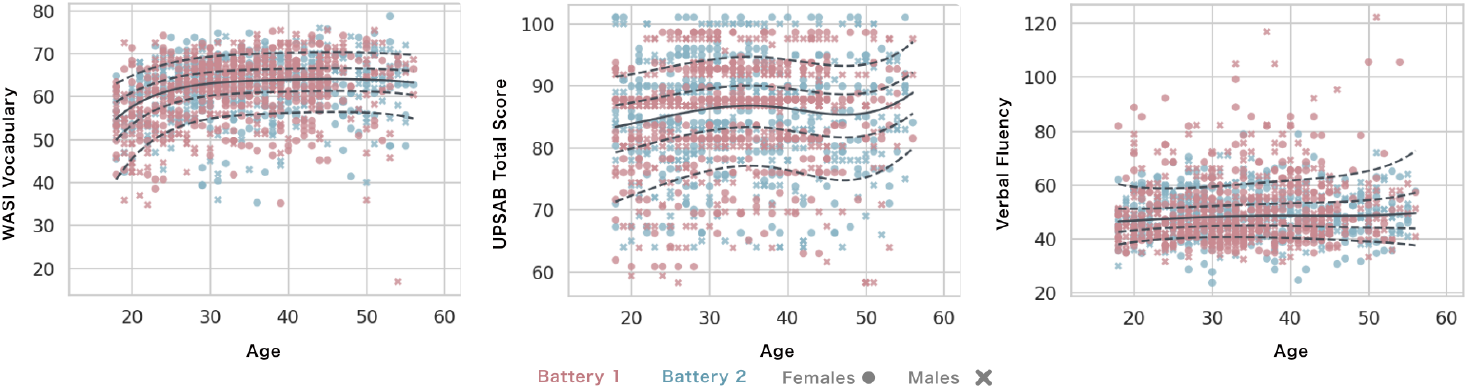
An example of normative models trained on the two imputed TOP batteries using the hierarchical method, depicted the variables shown in Fig. 6.

In summary, our findings highlight the varied impact of structured missingness on the performance of imputation methods. We show that varied evaluation metrics are needed to evaluate the effects of structured missingness. Further, we highlight that a narrow focus on numerical precision alone, risks obscuring underlying biases and site-specific variability, limiting the reliability of downstream analyses. By adopting approaches that preserve distributional integrity and multivariate relationships, we can better leverage heterogeneous, multi-site datasets to extract meaningful insights. This strategy enhances both subject-level inference and population-level modelling, as demonstrated by methods such as normative modelling. However, fully addressing the challenges posed by structured missingness requires the continued development of integrated imputation-harmonisation pipelines, improved fairness-aware validation frameworks, and adaptive modelling approaches capable of handling residual uncertainty. Ensuring robust and equitable imputation will be essential for advancing precision medicine and other areas of biomedical science.

## 4 Methods

Our simulation study comprised four distinct stages (Fig. 3). First, we synthesized complete datasets, thereby establishing a ground truth against which imputation accuracy could be assessed–a benchmark not generally in observational studies where true missing values remain unknown. Second, we introduced different patterns of missingness into the dataset, thereby creating an amputated dataset on which the imputation algorithms will be tested. We incorporated both random and structured missingness patterns to capture a range of real-world scenarios. Next, we applied the imputation algorithms to reconstruct the missing data. To account for the inherent uncertainty of these algorithms, each is initialised ten times using different random seeds, except for the MICE-based algorithm, which by defaults samples from matched donors for each variable without needing re-initialisation. Finally, the imputation results were evaluated using a comprehensive set of evaluation criteria. A detailed description of the process is outlined below.

### 4.1 Data simulation

The code used to generate the simulated dataset together with the imputation algorithms is available in a GitHub repository. The dataset simulation was performed as follows:

1. **Covariance structure:** A covariance matrix (Σ) and the mean vector (*µ*) were derived from a real dataset of cognition, ensuring realistic correlations between variables. This approach preserves the transitivity of the correlation matrix while adequately reflecting real-world scenarios. The covariance matrix was defined as:

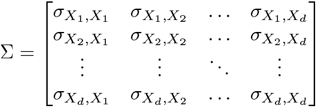

where each entry 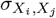 represents the estimated covariance between variables *X*_*i*_ and *X*_*j*_.
2. **Data generation:** Data were sampled from a multivariate normal distribution, with the mean vector and covariance matrix estimated from the cognitive dataset. To account for demographic effects, the data were conditioned on age (*X*_0_), which was generated uniformly between 18 and 60:

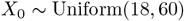 The relationship between age and each variable was modelled using linear regression:

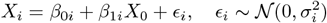

where *β*_0*i*_ and *β*_1*i*_ were estimated from the original cognitive dataset. Given a known value *X*_0_ = *x*_0_, the remaining variables were drawn from a conditional multivariate normal distribution:

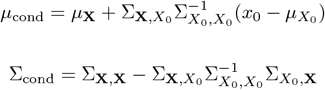

where 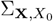 represents the covariance between *X*_0_ and the remaining variables **X** = (*X*_1_, …, *X*_*d*_).
3. **Site-specific effects:** To introduce variability across datasets, each variable’s mean 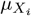 was randomly perturbed by a factor *δ*_*i*_:

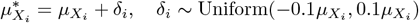
4. **Sample sizes:** Simulations were conducted with sample sizes of 100 and 1,000, split evenly between the two datasets. These sizes represent typical data scales in biomedical research, from exploratory studies to more extensive cohorts.

### 4.2 Missing data mechanisms

We introduced three forms of missingness, each simulated independently and in combination to reflect common challenges in dataset integration:

- **Random missingness:** For each variable *X*_*j*_, a predetermined percentage *p*_noise_ of values was set as missing at random. Missingness probability for each variable was applied uniformly across observations:

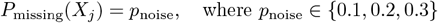 This approach reflects the missing at random with no structure mechanism.
- **Blockwise missingness across datasets:** Variables were randomly assigned to one of two datasets, *D*_1_ or *D*_2_, while maintaining a predefined overlap set of variables. The overlap size was set to 4, 10, or 20 variables to reflect low, medium, and high overlap scenarios. Variables outside the overlap set were deleted from the alternate dataset to create a blockwise missing structure:

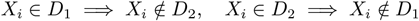 This approach simulates the multi-dataset contexts with partially overlapping variables.
- **Dependency missingness:** Following **[16]**, dependencies in missingness were introduced between *two* selected columns. The pattern was implemented as follows:
  1. **Predictive variable standardisation:** For a predictor variable *X*_*i*_ shared across datasets, we standardised it by computing:

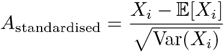
  2. **Probability of missingness:** Missingness in a dependent variable *Y* was assigned based on the standardised predictor’s deviation, using:

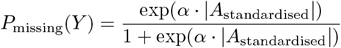

where *α* = 3 controls the likelihood of missingness based on the predictor’s distribution characteristics.
  3. **Conditional missingness in additional variables:** For a third variable *Z*, missingness was introduced conditionally on *Y* ‘s missing status:

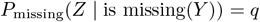

with *q* = *missingness*, specifying a structured, conditional missingness pattern.

### 4.3 Imputation Methods

#### 4.3.1 Average Imputation

Average imputation is the most straightforward imputation method, where the mean of each variable is used to replace all the missing values for that variable. While this method is not advocated for practical use, it serves as a demonstration of its impact on data distribution and other quality measures. Additionally, it provides a basic benchmark against which the performance of more advanced imputation algorithms can be evaluated.

#### 4.3.2 Extremely Randomised Trees

Extremely Randomised Trees (Extra Trees) is a tree-based ensemble method for classification and regression that introduces additional randomness during tree construction by randomising both the feature selection and split threshold choice **[36]**. Unlike Random Forests, which select the best split from a random subset of features, Extra Trees chooses the feature and the cut-point completely at random, promoting greater diversity among the trees in the ensemble and improving robustness in some cases **[36, 51]**. Furthermore, unlike in the Random Forest case, each decision tree in the ensemble is built on the entire dataset (without bootstrapping). The results are then derived by averaging (regression) or majority voting (classification). We have chosen this method, as it has been previously shown to adapt well to nonlinearities in datasets, often a desirable property for imputation in complex datasets **[6, 52]**. We took advantage of the python implementation of random forests, combined with the Iterative Imputer implementation for imputation.

#### 4.3.3 AutoComplete

AutoComplete is a recently introduced imputation algorithm aimed at handling extensive phenotypic missingness in large-scale biobank datasets **[30]**. Using a copy-masking mechanism within an autoencoder architecture, it learns patterns within incomplete data to impute both binary and continuous phenotypes. Autoencoders are a type of neural network consisting of an encoder that compresses data into a low-dimensional latent layer, followed by a decoder that reconstructs the data from this compressed representation. Copy-masking allows the algorithm to incorporate partially observed data by selectively masking inputs, effectively increasing sample-size suitable for training as well as improving imputation robustness. While AutoComplete demonstrated improved performance over traditional imputation methods, it has not yet been evaluated on datasets with block-structured missingness, leaving open questions about its effectiveness with structured missingness scenarios, a common feature in integrated biobank data.

#### 4.3.4 Multivariate Imputation by Chained Equations (MICE)

The Fully Conditional Specification approach, commonly implemented as Multivariate Imputation by Chained Equations **[31]**, is a widely adopted framework for multiple imputation. MICE operates iteratively, specifying a conditional model for each variable given all others, and then draws imputed values from the (approximate) posterior predictive distribution of these models. This inherent stochasticity, rather than imputing a single best prediction, introduces appropriate random variation, crucial for preserving data variance and reflecting imputation uncertainty. In this study, we utilised Predictive Mean Matching within the MICE framework. This algorithm first fits a temporary regression model using observed data to predict values for missing entries. It then identifies a small set of donor cases from the observed data whose predicted values are closest to that of the missing entry. Finally, an actual observed value is randomly selected from one of these donors as the imputation, thereby ensuring that imputed values are plausible and drawn from the observed data distribution.

#### 4.3.5 Hierarchical MICE with Site-Specific Imputation

To account for multi-site variability, we implemented a hierarchical extension of the Multivariate Imputation by Chained Equations (MICE) **[32]**, explicitly modelling site-specific offsets for variables measured across multiple locations. This approach ensures that local differences are preserved while maintaining consistency across the dataset.

Variables were first classified based on their site-specific missingness patterns. Those measured across multiple sites were flagged for hierarchical imputation, while fully observed variables remained unchanged. A *prediction matrix* was constructed using pairwise correlations, retaining only variable pairs with sufficient non-missing observations. Site-specific effects were incorporated by introducing an offset for multi-site variables, modelled as:

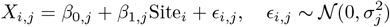

where *β*_1*j*_ captures site-level deviations. Single-site variables were imputed without this adjustment. Such strategy balances local variability with overall coherence, refining imputation accuracy in multi-site studies.

### 4.4 Evaluation Metrics

The following metrics have been used for evaluation of simulations, where the ground truth is available (Fig. 3).

#### 4.4.1 Coverage and Width

Coverage and width are metrics used to quantify the quality of imputation at the subject level. Coverage represents the proportion of 95% confidence intervals across all subjects that include the true value. Ideally, 100% of the intervals would contain the true value. However, this condition can be trivially achieved by making the intervals infinitely wide. Wider intervals increase the likelihood of containing the true value but provide little to no meaningful information for individual subjects. To balance this, the width of the 95% confidence intervals is also a critical measure. The intervals should be as narrow as possible while maintaining adequate coverage, ensuring they provide precise and informative estimates.

#### 4.4.2 Mean squared error and distribution fit

For summarizing imputation performance for individual variables, we use the mean squared error, but normalise it by the variance of each variable to avoid the disproportionate influence of variables with larger numerical ranges. Additionally, instead of reporting the normalised mean squared error (NMSE) directly, we report 1 *− NMSE* for easier interpretation. In this scale, a value of 1 indicates perfect imputation, a value of 0 corresponds to performance equal to imputing with the variable’s mean, and values below 0 indicate worse-than-average imputation.

While NMSE effectively measures the accuracy of imputed values, it does not assess whether the original distribution of the data has been preserved. To address this, we use the Jensen–Shannon divergence (JSD; equation 1). JSD is a variation of the more widely known Kullback–Leibler divergence but has the advantages of being symmetric and bounded between 0 and 1. These properties make it more suitable for comparing variables with different scales. In our formulation, *p* and *q* denote the probability distributions of the observed and imputed data, respectively, and 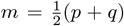 is their average. For consistency, we report 1 − JSD, where a value of 1 indicates perfect alignment between the original and imputed distributions, and a value of 0 indicates completely dissimilar distributions.

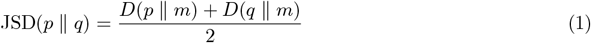

#### 4.4.3 Multivariate relationships

When imputed variables are intended for use in machine learning, preserving multivariate relationships becomes critically important. The quality of imputation has been shown to significantly influence second-stage machine learning outcomes and algorithmic fairness **[35, 53]**. However, the preservation of these complex relationships is often overlooked in imputation evaluations.

To address this, we define two metrics to assess multivariate relationships. The first is the Frobenius norm, calculated as the difference between two correlation matrices, the original and imputed. Since correlation matrices are symmetrical, we focus on subtracting the upper triangle of one matrix from the other. The second metric is the correlation of the vectorised upper triangles of the two matrices. The Frobenius norm provides a summary measure of how well the overall correlations are preserved during imputation, while the ‘correlation of correlations’ captures whether the relationships between variables are retained or if systematic distortions emerge.

## 5 Data

To evaluate imputation algorithms in a real world scenario, we analysed cognitive battery data from the healthy subjects of the Norwegian ‘Thematically Organised Psychosis’ (TOP) cohort (Table 2) **[37, 54]**. This dataset offers a controlled setting where common sources of structured missingness, such as variations in language, personnel, and testing location, are minimised. Instead, missingness arose specifically from a transition between two cognitive test batteries (Battery 1, 2004-2012 and Battery 2, 2012-2023), which were designed to measure similar constructs but used different tests. The dataset comprised 22 overlapping test scores, with the first battery including 11 additional unique scores and the second containing 5 unique scores. A complete list of test scores is provided in Appendix 2.

**Table 2:** Demographics of the samples stratified by datasets.

|  | Site | Sex | count | Age (min, max) |
| --- | --- | --- | --- | --- |
| TOP | 1 | All | 509 | 34.6 (18, 56) |
|  |  | Females | 254 | 34.7 (18, 55) |
|  |  | Males | 255 | 34.4 (18, 56) |
|  | 2 | All | 667 | 33.6 (18, 56) |
|  |  | Females | 306 | 33.3 (18, 56) |
|  |  | Males | 361 | 33.9 (18, 56) |
| CNSR | 3 | All | 80 | 26.9 (18, 39) |
|  |  | Females | 25 | 26.9 (18, 38) |
|  |  | Males | 55 | 27 (20, 39) |
|  | 4 | All | 119 | 23.6 (18, 42) |
|  |  | Females | 61 | 22.1 (18, 34) |
|  |  | Males | 58 | 25.2 (19, 42) |
|  | 5 | All | 100 | 23.5 (18, 38) |
|  |  | Females | 56 | 22.3 (18, 33) |
|  |  | Males | 44 | 25 (19, 38) |

To further assess imputation performance across diverse settings, we incorporated additional healthy cohorts from the Centre for Neuropsychiatric Schizophrenia Research in Denmark (Table 2) **[38–40]**. These datasets, collected as part of distinct studies, exhibited substantial but not complete overlap in measured variables. Fig. 8 details the number of overlapping and unique test scores in all centres, while the demographic characteristics are presented in Table 2. The inclusion of additional datasets introduced further challenges for imputation. First, site effects were expected to be more pronounced due to differences in language (which required test translation), testing personnel, and cohort characteristics. Simultaneously, the expanded dataset increased overall variability, potentially benefiting downstream analyses by improving generalisability. The imputation was performed using all algorithms tested in the simulation studies and each algorithm was run ten times to assess the variability and uncertainty.

**Figure 8:**
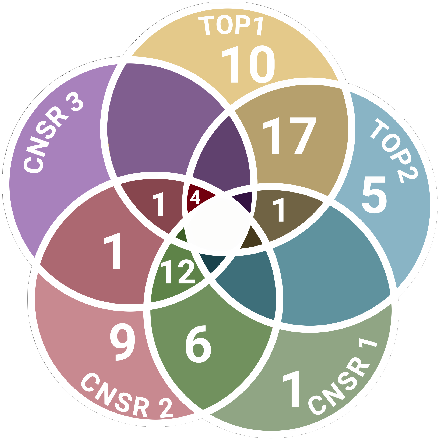
Overlap of conducted tests across the evaluated datasets.

Two complementary evaluation strategies were used. First, since real-world imputation lacks a direct ground truth, we leveraged observed values through k-fold cross-validation, following the approach of Llera et al. **[6]**. In each fold, 5% of observed values were randomly removed in addition to the already missing values, and the full dataset was subsequently imputed **[6]**. This procedure generated reconstructed values for originally measured variables, allowing us to assess imputation accuracy using the metrics defined above. While this provides the closest possible approximation to a ground-truth evaluation, it is important to note that this approach does not directly capture the challenges introduced by structured missingness, as there is no ground truth for systematically missing values.

To specifically address structured missingness, we employed JSD to quantify the extent to which the distribution of imputed values aligned with that of the observed values. For each variable, we compared the distribution of measured values with that of the imputed values. If no values were missing for a given variable, the JSD was not estimated. For improved clarity and readability, all results have been concatenated and presented in the form of Figures in the Results section (Fig. 4) and Appendix 1.

## 7 Appendix 1

Here, we present additional results from the simulation study, specifically examining the effects of varying levels of added random noise, both in the context of random missingness and when combined with block and dependence missingness. The purpose of this extension is to assess the robustness of imputation algorithms under different noise conditions, simulating real-world variability in multi-site datasets.

We consider scenarios with overlapping variable sets of sizes 4, 10, and 20, introducing random noise to 10%, 30%, and 50% of values. This noise affects only the randomly missing and dependent missing values but does not impact block missingness, which remains at 100% for a given site by definition.

**Figure 9:**
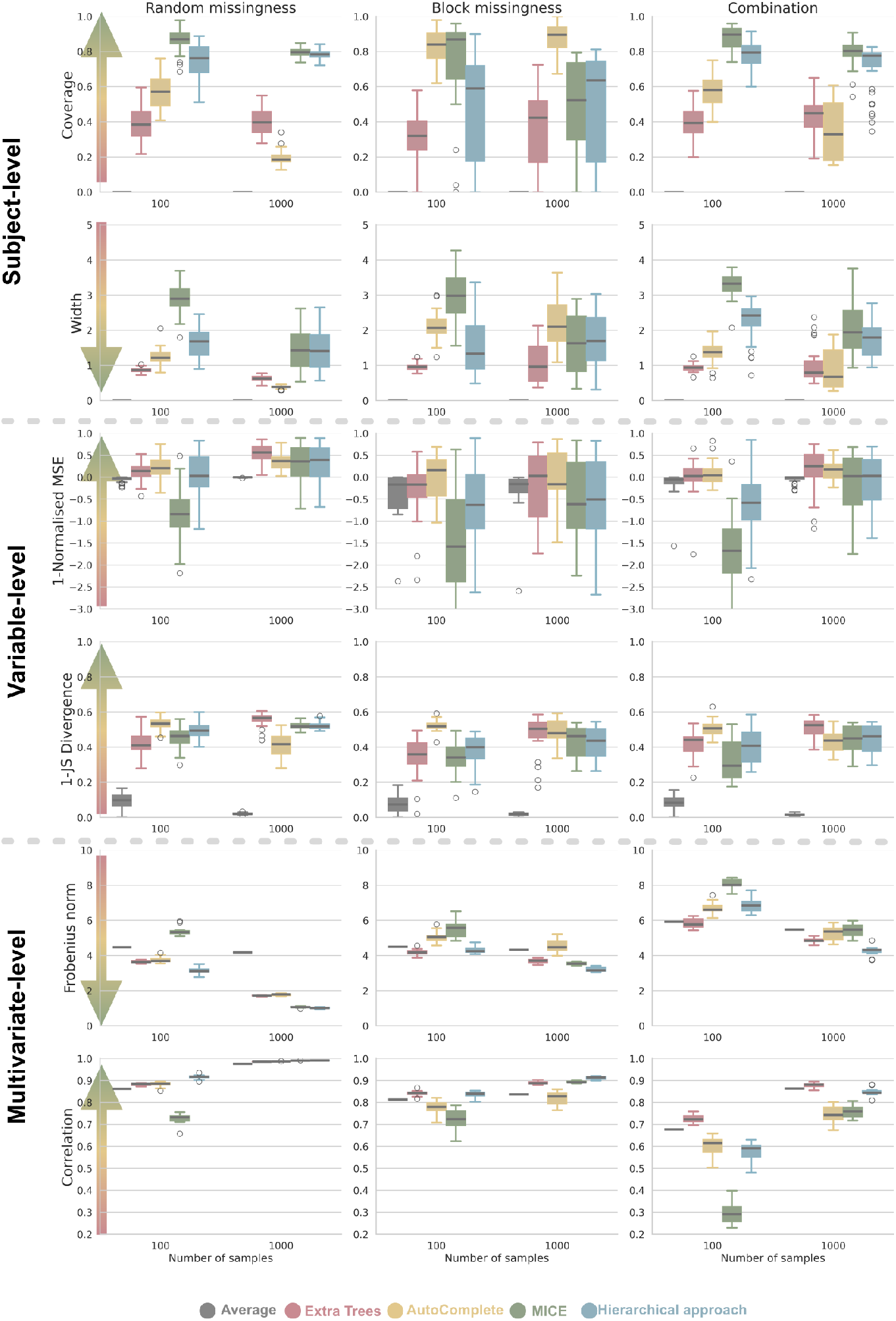
Evaluation of the imputation methods on the simulated datasets: Each row represents a quality of imputation criterion, and each column represents a different type of missingness. The results are shown for five imputation algorithms: Average imputation, Extra Trees, AutoComplete, MICE, and hierarchical MICE; and for two different dataset sizes (100, 1000). Here, the amount of random noise missingness was set to 50%, while both batteries overlapped in 20 variables. The arrows on the left side point in the direction of the ideal reconstruction values.

**Figure 10:**
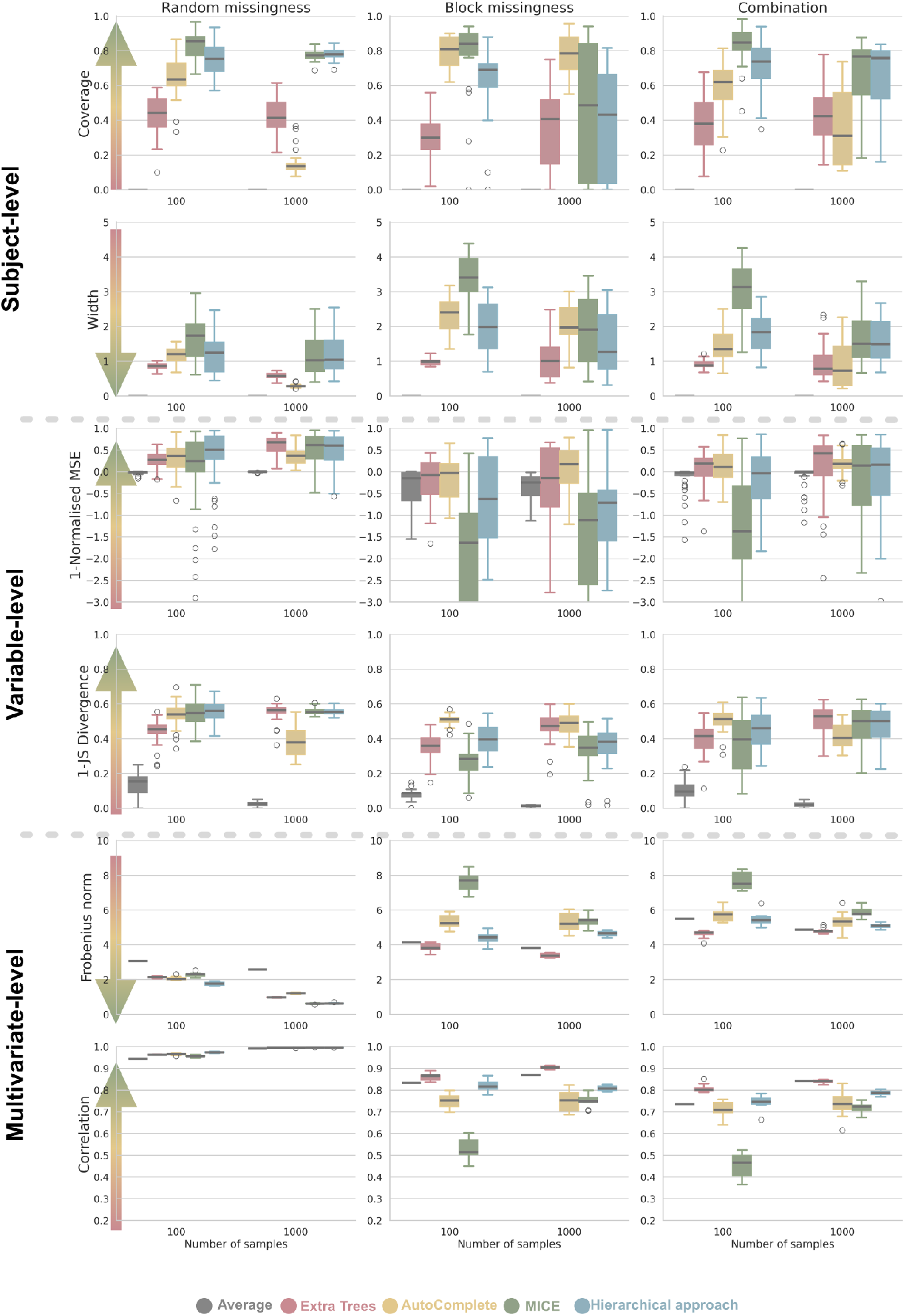
Evaluation of the imputation methods on the simulated datasets: Each row represents a quality of imputation criterion, and each column represents a different type of missingness. The results are shown for five imputation algorithms: Average imputation, Extra Trees, AutoComplete, MICE, and hierarchical MICE; and for two different dataset sizes (100, 1000). Here, the amount of random noise missingness was set to 10%, while both batteries overlapped in 20 variables. The arrows on the left side point in the direction of the ideal reconstruction values.

**Figure 11:**
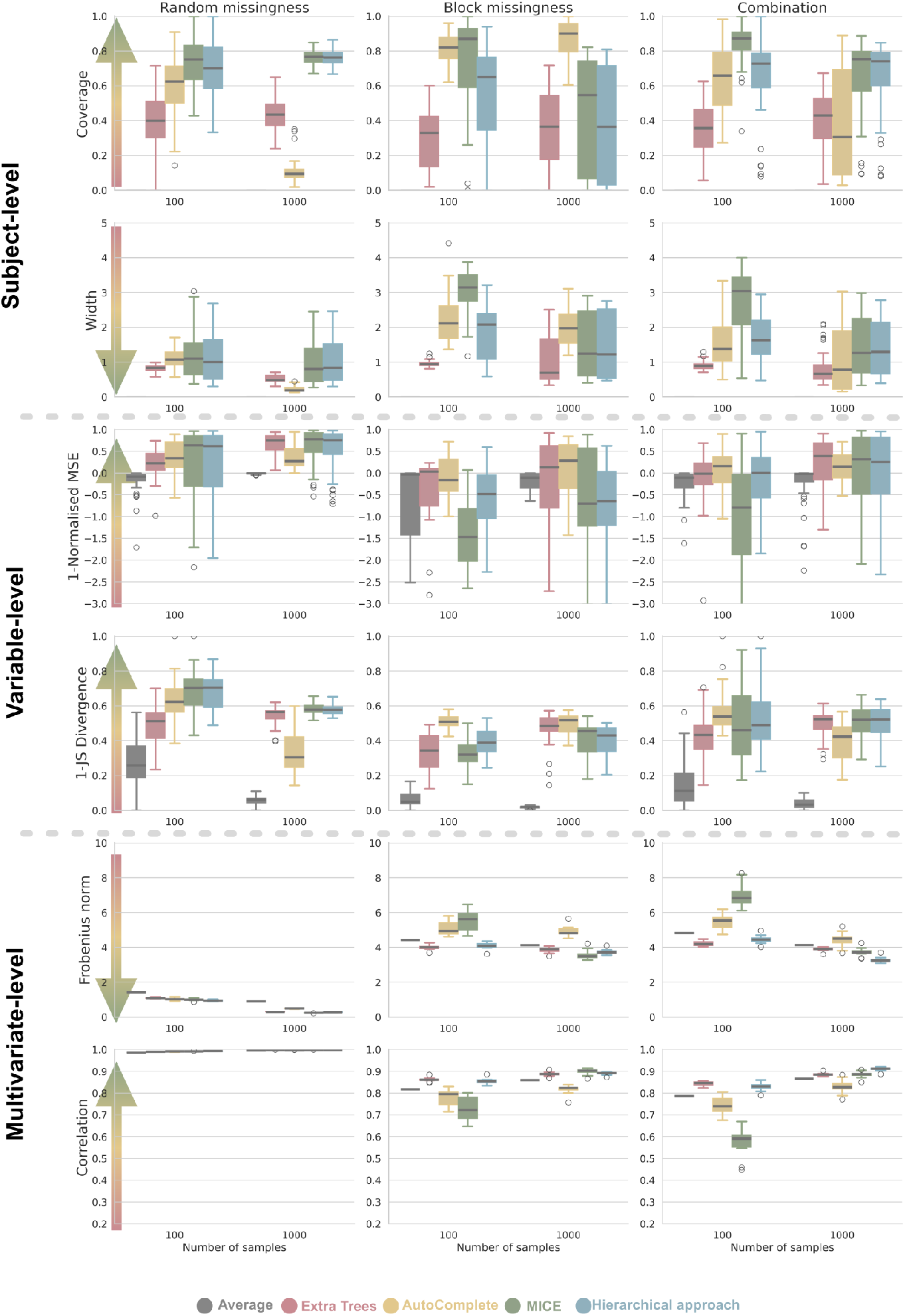
Evaluation of the imputation methods on the simulated datasets: Each row represents a quality of imputation criterion, and each column represents a different type of missingness. The results are shown for five imputation algorithms: Average imputation, Extra Trees, AutoComplete, MICE, and hierarchical MICE; and for two different dataset sizes (100, 1000). Here, the amount of random noise missingness was set to 10%, while both batteries overlapped in 20 variables. The arrows on the left side point in the direction of the ideal reconstruction values.

**Figure 12:**
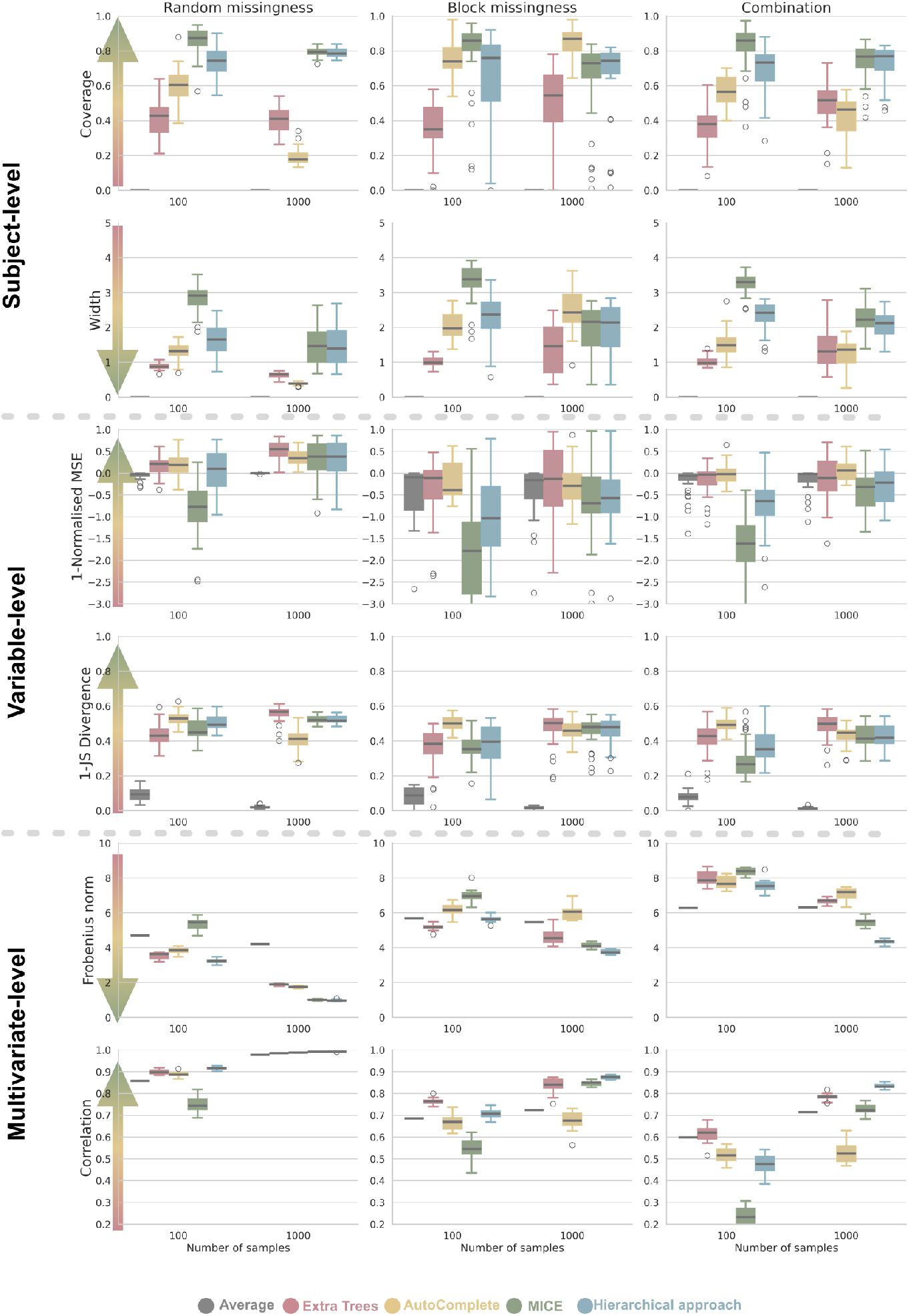
Evaluation of the imputation methods on the simulated datasets: Each row represents a quality of imputation criterion, and each column represents a different type of missingness. The results are shown for five imputation algorithms: Average imputation, Extra Trees, AutoComplete, MICE, and hierarchical MICE; and for two different dataset sizes (100, 1000). Here, the amount of random noise missingness was set to 50%, while both batteries overlapped in 10 variables. The arrows on the left side point in the direction of the ideal reconstruction values.

**Figure 13:**
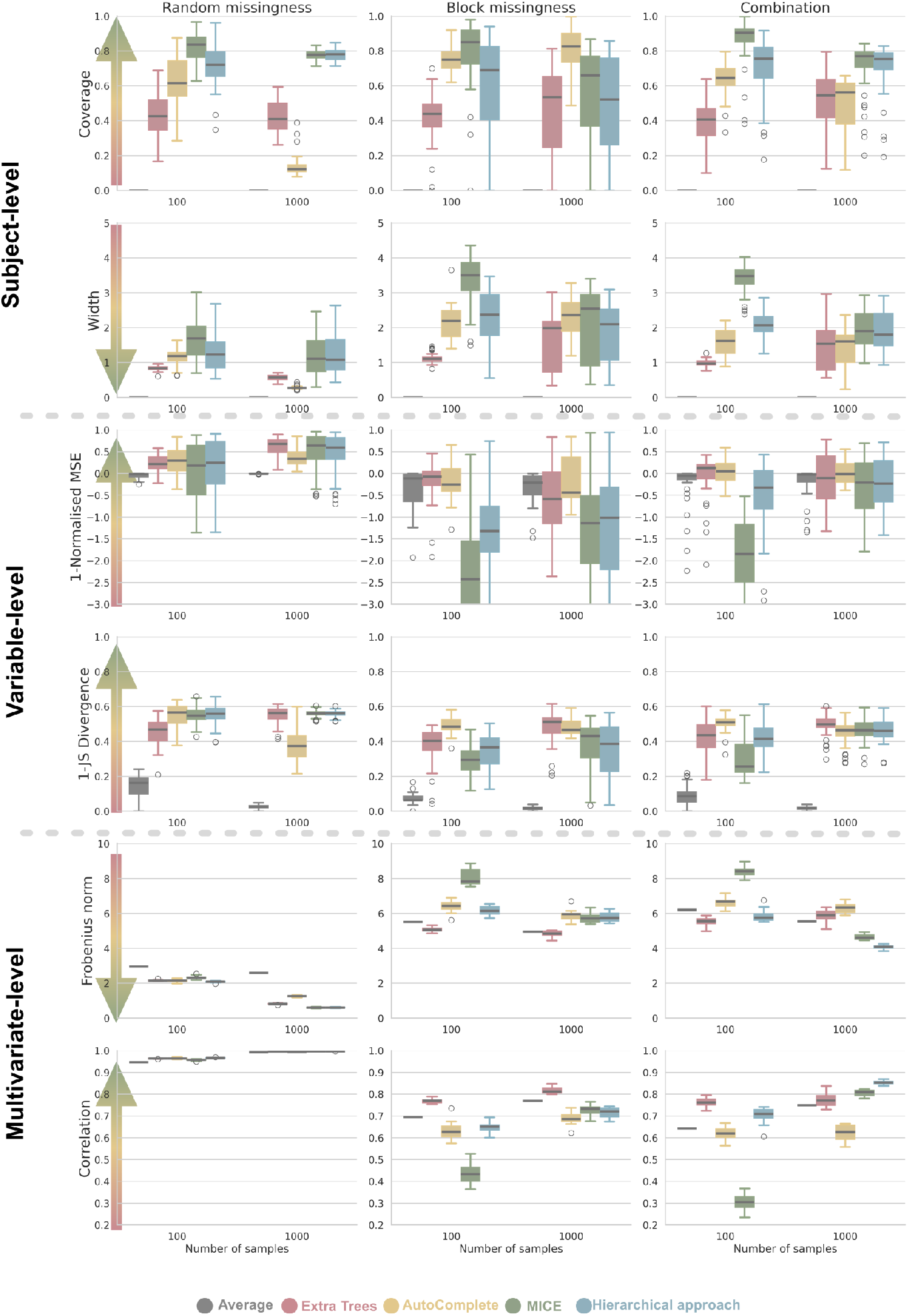
Evaluation of the imputation methods on the simulated datasets: Each row represents a quality of imputation criterion, and each column represents a different type of missingness. The results are shown for five imputation algorithms: Average imputation, Extra Trees, AutoComplete, MICE, and hierarchical MICE; and for two different dataset sizes (100, 1000). Here, the amount of random noise missingness was set to 30%, while both batteries overlapped in 10 variables. The arrows on the left side point in the direction of the ideal reconstruction values.

**Figure 14:**
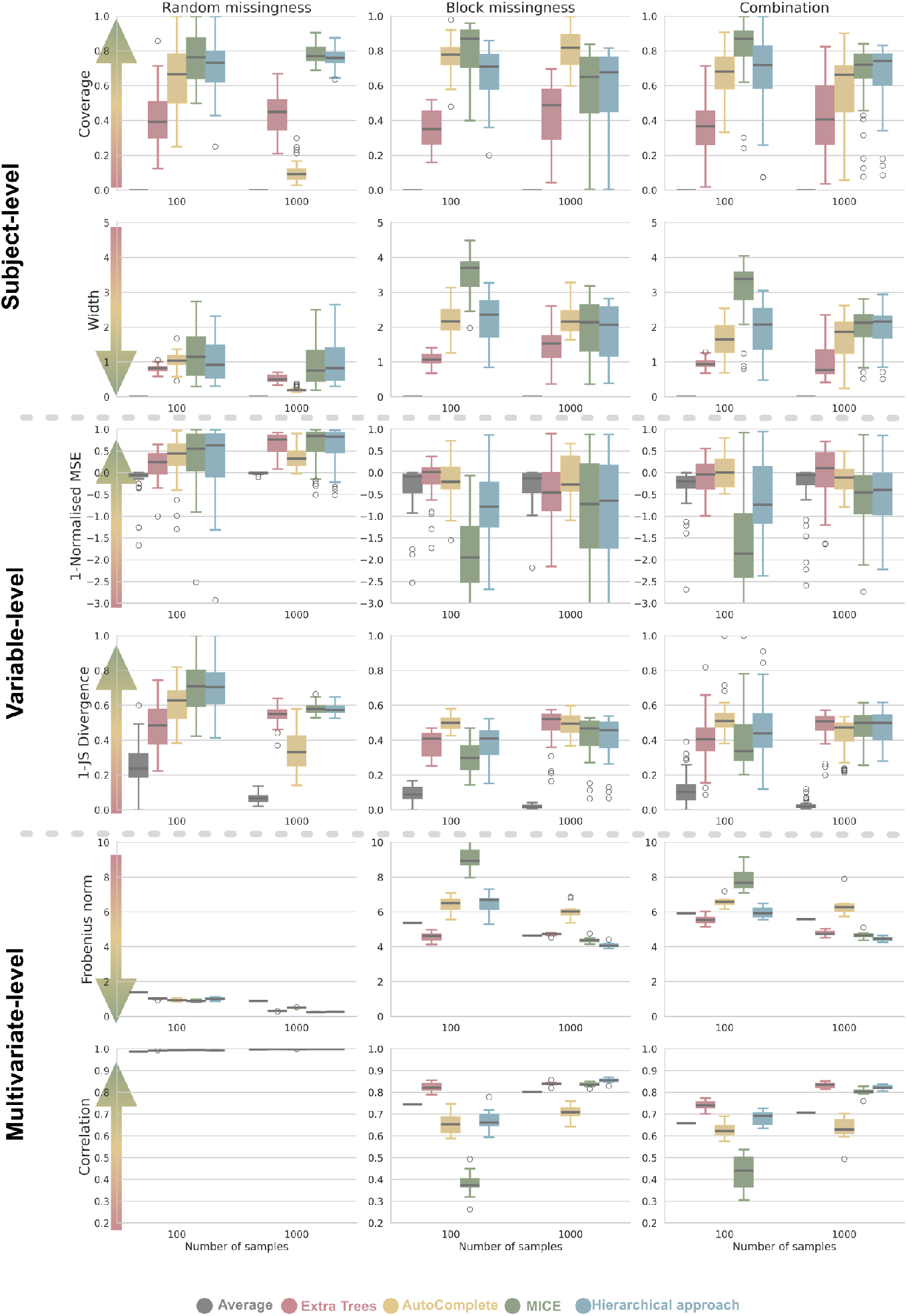
Evaluation of the imputation methods on the simulated datasets: Each row represents a quality of imputation criterion, and each column represents a different type of missingness. The results are shown for five imputation algorithms: Average imputation, Extra Trees, AutoComplete, MICE, and hierarchical MICE; and for two different dataset sizes (100, 1000). Here, the amount of random noise missingness was set to 10%, while both batteries overlapped in 10 variables. The arrows on the left side point in the direction of the ideal reconstruction values.

**Figure 15:**
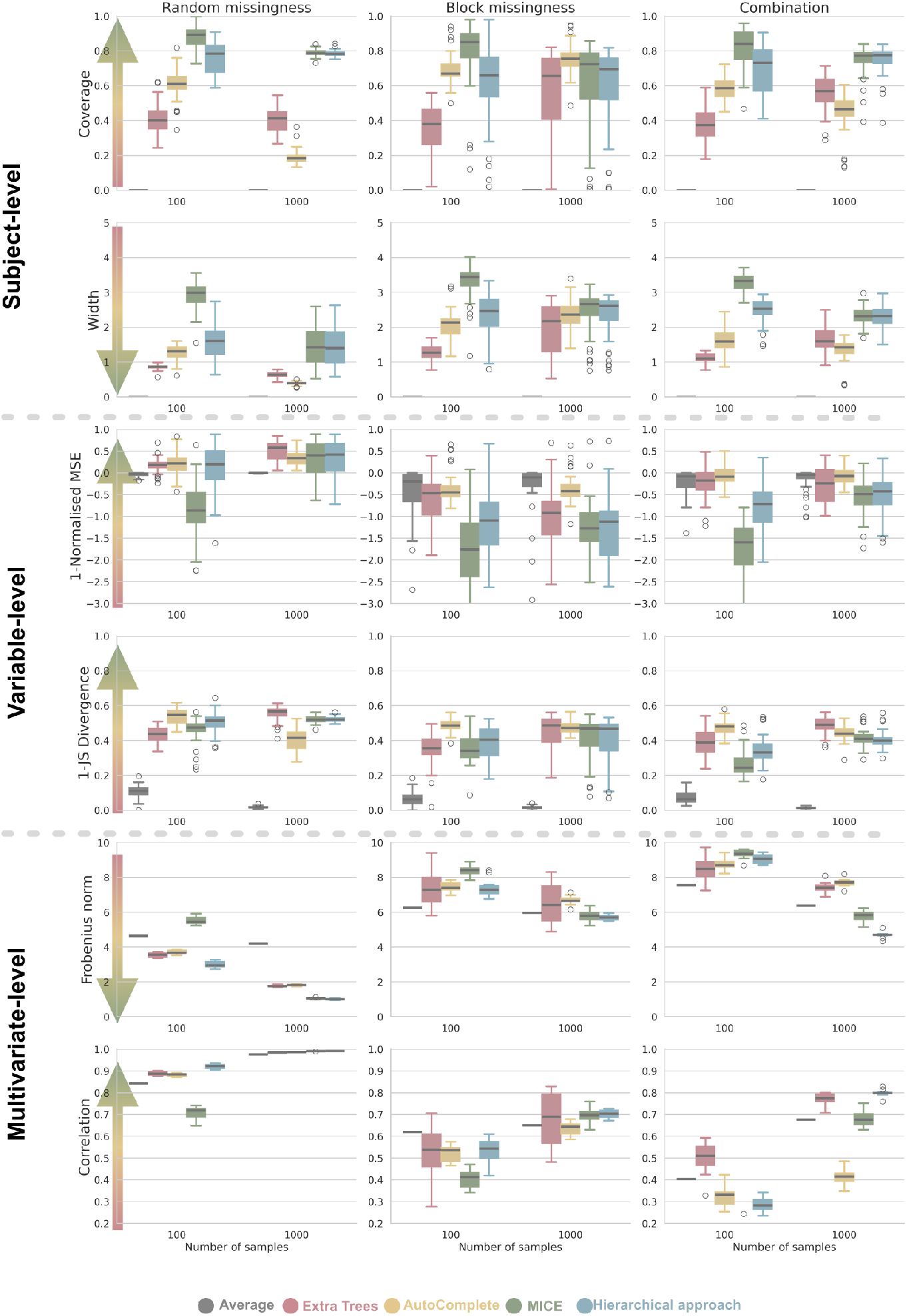
Evaluation of the imputation methods on the simulated datasets: Each row represents a quality of imputation criterion, and each column represents a different type of missingness. The results are shown for five imputation algorithms: Average imputation, Extra Trees, AutoComplete, MICE, and hierarchical MICE; and for two different dataset sizes (100, 1000). Here, the amount of random noise missingness was set to 50%, while both batteries overlapped in 4 variables. The arrows on the left side point in the direction of the ideal reconstruction values.

**Figure 16:**
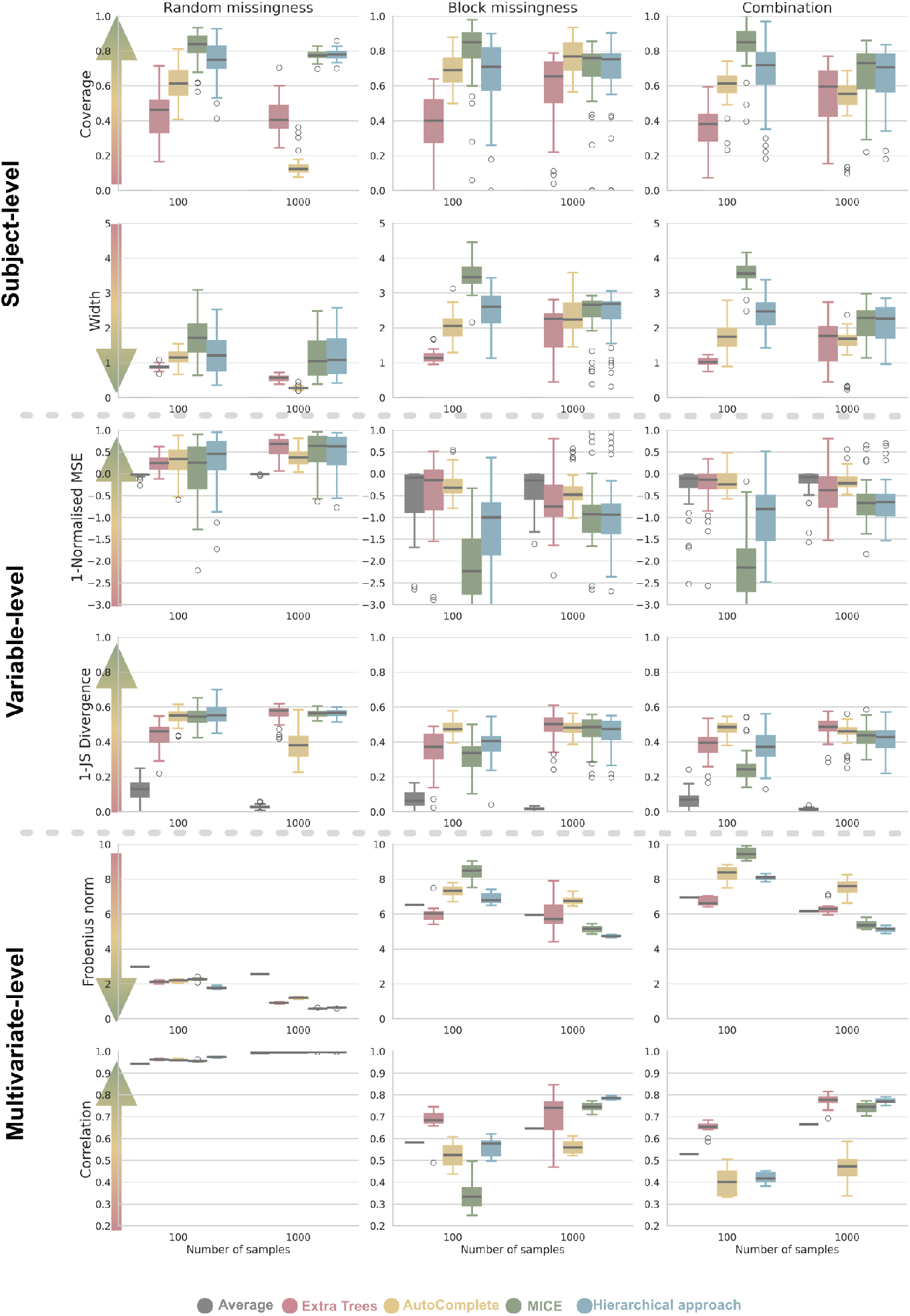
Evaluation of the imputation methods on the simulated datasets: Each row represents a quality of imputation criterion, and each column represents a different type of missingness. The results are shown for five imputation algorithms: Average imputation, Extra Trees, AutoComplete, MICE, and hierarchical MICE; and for two different dataset sizes (100, 1000). Here, the amount of random noise missingness was set to 30%, while both batteries overlapped in 4 variables. The arrows on the left side point in the direction of the ideal reconstruction values.

**Figure 17:**
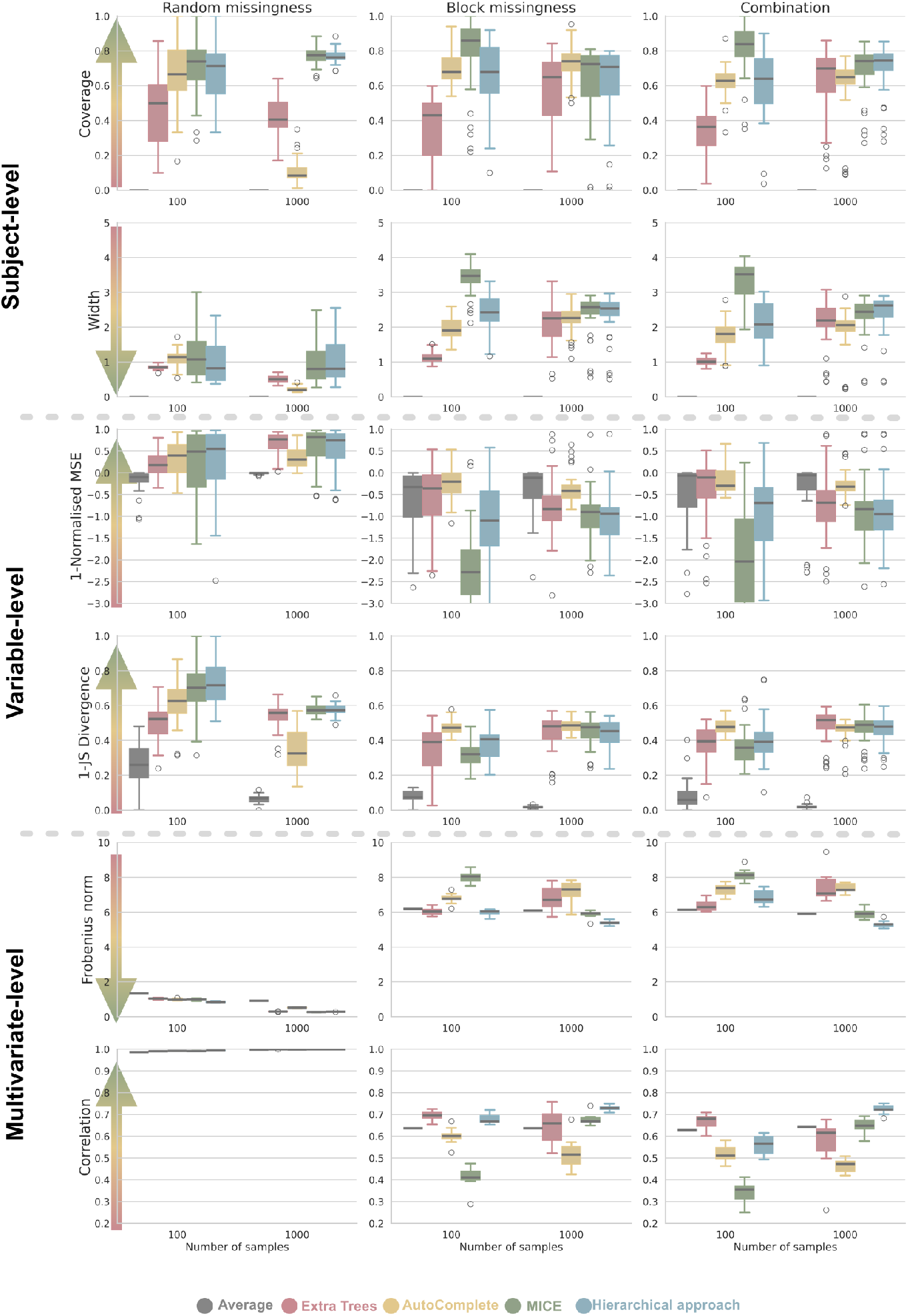
Evaluation of the imputation methods on the simulated datasets: Each row represents a quality of imputation criterion, and each column represents a different type of missingness. The results are shown for five imputation algorithms: Average imputation, Extra Trees, AutoComplete, MICE, and hierarchical MICE; and for two different dataset sizes (100, 1000). Here, the amount of random noise missingness was set to 10%, while both batteries overlapped in 4 variables. The arrows on the left side point in the direction of the ideal reconstruction values.

## 8 Appendix 2

Below, we present a table of the cognitive tests measured in all seven tested cohorts.

**Table 3:**
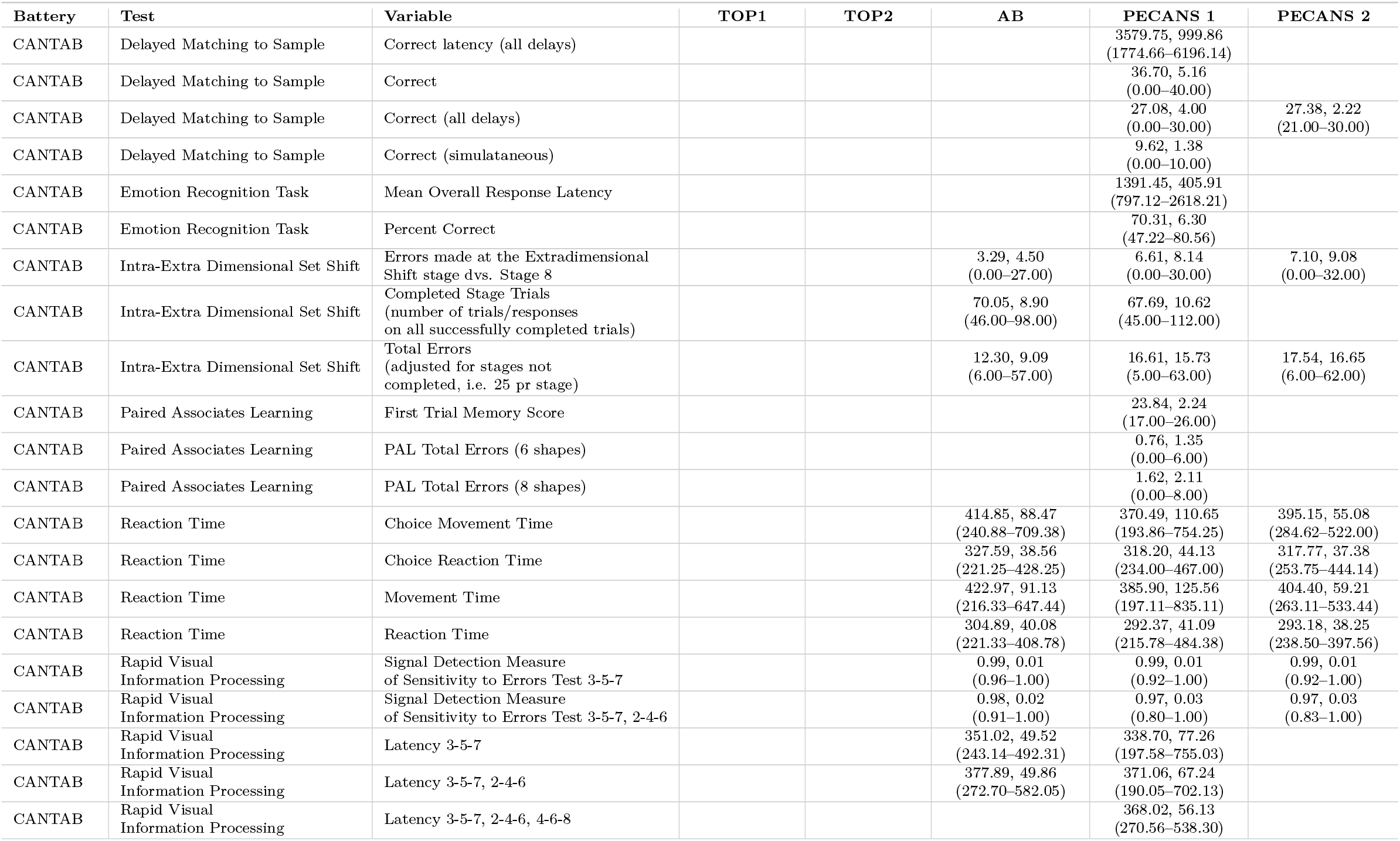
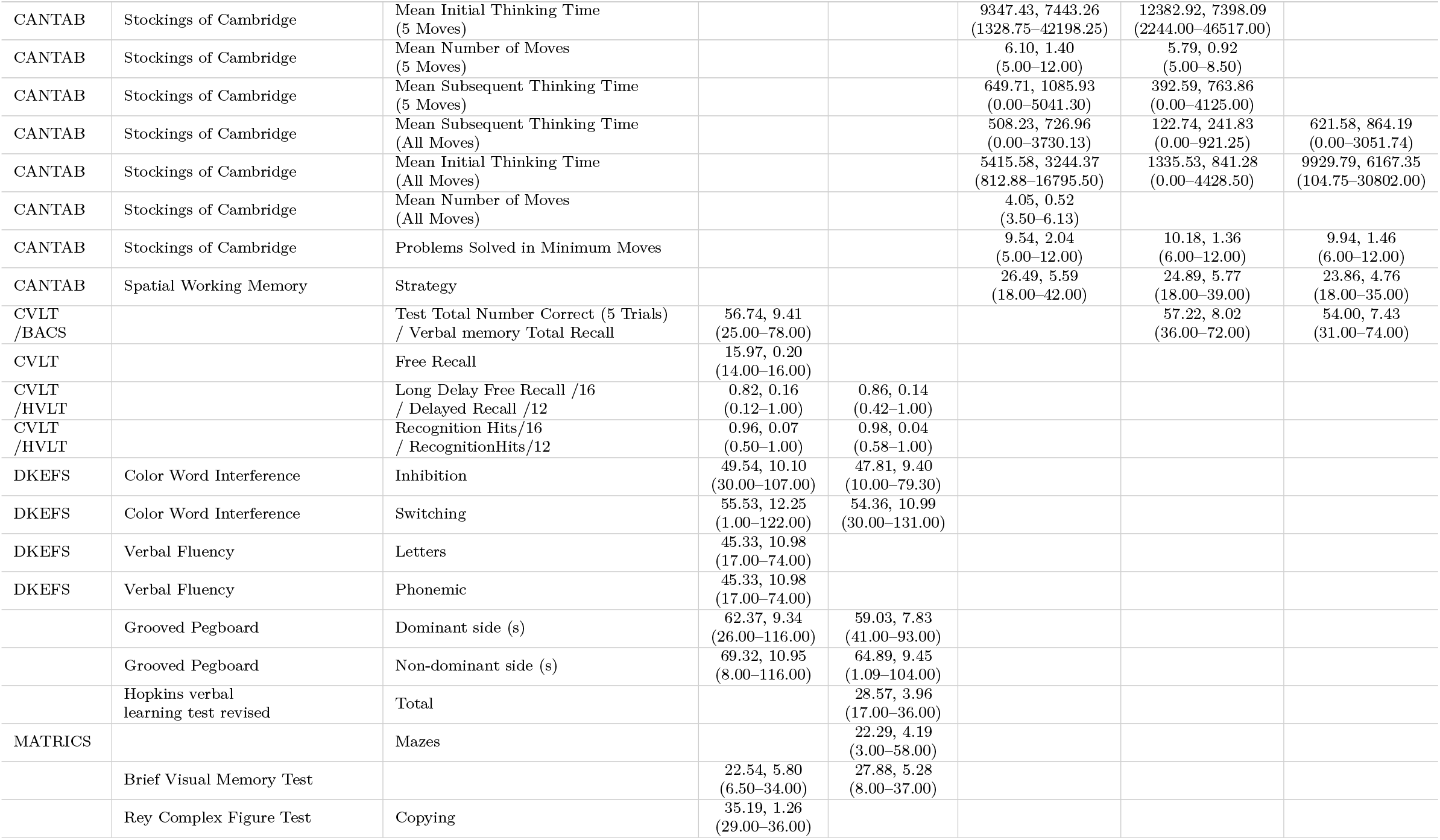
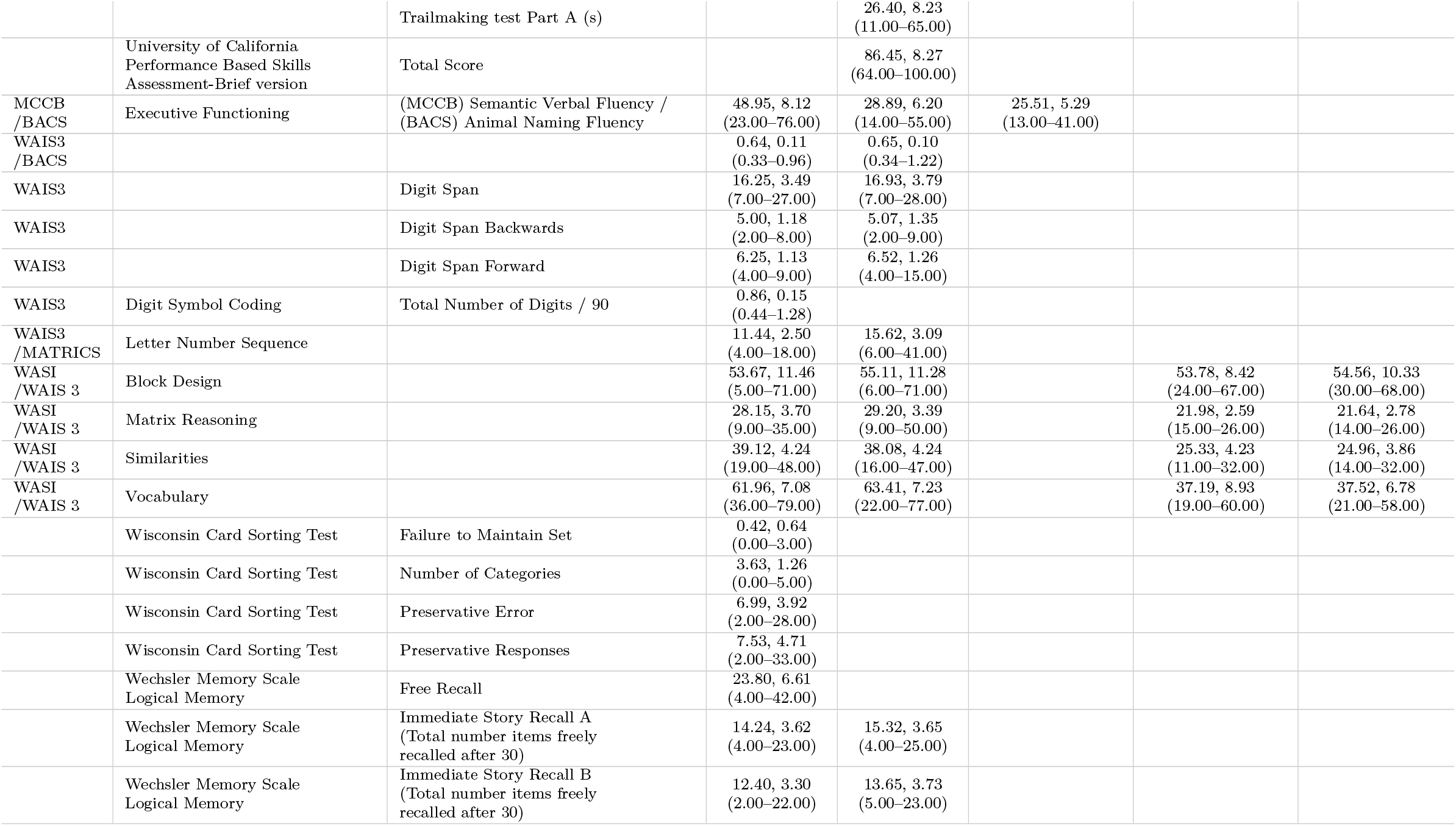
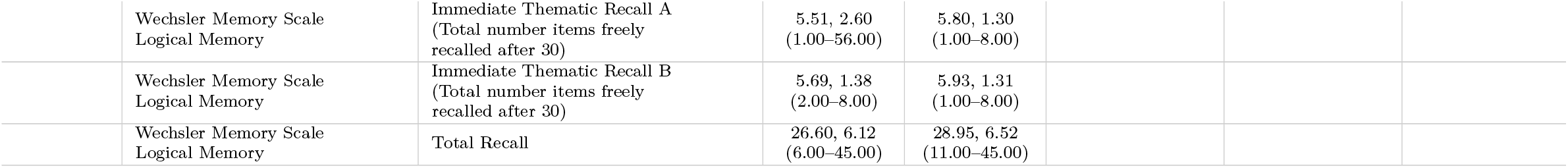
Descriptive statistics (mean, standard deviation, and range) for cognitive test variables across datasets: TOP1, TOP2, AB, PECANS1, and PECANS2. Variables reflect various cognitive domains such as memory, attention, emotion recognition, executive function, and processing speed. Empty cells indicate missing data for that particular dataset and variable.

## References

1. Rutherford, S. et al. Charting Brain Growth and Aging at High Spatial Precision. eLife 11 (eds Baker, C. I., Taschler, B., Esteban, O. & Constable, T.) e72904. ISSN: 2050-084X. 10.7554/eLife.72904 (2024) (Feb. 1, 2022).

2. Bethlehem, R. a. I. et al. Brain Charts for the Human Lifespan. Nature 604, 525–533. ISSN: 1476-4687. https://www.nature.com/articles/s41586-022-04554-y (2024) (Apr. 2022).

3. Kennedy, E. et al. Bridging Big Data: Procedures for Combining Non-equivalent Cognitive Measures from the ENIGMA Consortium https://www.biorxiv.org/content/10.1101/2023.01.16.524331v2 (2024). Pre-published.

4. Li, J. et al. Imputation of Missing Values for Electronic Health Record Laboratory Data. npj Digital Medicine 4, 1–14. ISSN: 2398-6352. https://www.nature.com/articles/s41746-021-00518-0 (2024) (Oct. 11, 2021).

5. Sousa da Mota, B. et al. Imputation of Ancient Human Genomes. Nature Communications 14, 3660. ISSN: 2041-1723. https://www.nature.com/articles/s41467-023-39202-0 (2024) (June 20, 2023).

6. Llera, A. et al. Evaluation of Data Imputation Strategies in Complex, Deeply-Phenotyped Data Sets: The Case of the EU-AIMS Longitudinal European Autism Project. BMC medical research methodology 22, 229. ISSN: 1471-2288. PMID: 35971088 (Aug. 16, 2022).

7. Mitra, R. et al. Learning from Data with Structured Missingness. Nature Machine Intelligence 5, 13–23. ISSN: 2522-5839. https://www.nature.com/articles/s42256-022-00596-z (2024) (Jan. 2023).

8. Rubin, D. B. Inference and Missing Data (1976).

9. Rubin, D. B. Formalizing Subjective Notions about the Effect of Nonrespondents in Sample Surveys. Journal of the American Statistical Association 72, 538–543. ISSN: 0162-1459. https://www.tandfonline.com/doi/abs/10.1080/01621459.1977.10480610 (2024) (Sept. 1, 1977).

10. Rubin, D. B. Multiple Imputation after 18+ Years. Journal of the American Statistical Association 91, 473–489. ISSN: 0162-1459, 1537-274X. http://www.tandfonline.com/doi/abs/10.1080/01621459.1996.10476908 (2024) (June 1996).

11. Enders, C. K. A Primer on Maximum Likelihood Algorithms Available for Use With Missing Data. Structural Equation Modeling: A Multidisciplinary Journal 8, 128–141. ISSN: 1070-5511, 1532-8007. http://www.tandfonline.com/doi/abs/10.1207/S15328007SEM0801_7 (2024) (Jan. 2001).

12. Enders, C. K. & Bandalos, D. L. The Relative Performance of Full Information Maximum Likelihood Estimation for Missing Data in Structural Equation Models. Structural Equation Modeling: A Multidisciplinary Journal 8, 430–457. ISSN: 1070-5511. https://www.tandfonline.com/doi/abs/10.1207/S15328007SEM0803_5 (2024) (July 1, 2001).

13. Lee, T. & Shi, D. A Comparison of Full Information Maximum Likelihood and Multiple Imputation in Structural Equation Modeling with Missing Data. Psychological Methods 26, 466–485. ISSN: 1939-1463, 1082-989X. http://doi.apa.org/getdoi.cfm?doi=10.1037/met0000381 (2024) (Aug. 2021).

14. Little, R. J., Rubin, D. B. & Zangeneh, S. Z. Conditions for Ignoring the Missing-Data Mechanism in Likelihood Inferences for Parameter Subsets. Journal of the American Statistical Association 112, 314–320. ISSN: 0162-1459. 10.1080/01621459.2015.1136826 (2024) (Jan. 2, 2017).

15. Jakobsen, J. C., Gluud, C., Wetterslev, J. & Winkel, P. When and How Should Multiple Imputation Be Used for Handling Missing Data in Randomised Clinical Trials – a Practical Guide with Flowcharts. BMC Medical Research Methodology 17, 162. ISSN: 1471-2288. 10.1186/s12874-017-0442-1 (2024) (Dec. 6, 2017).

16. Jackson, J., Mitra, R., Hagenbuch, N., McGough, S. & Harbron, C. A Complete Characterisation of Structured Missingness 2307.02650 [stat]. http://arxiv.org/abs/2307.02650 (2024). Pre-published.

17. Li, J. & Lomax, R. G. Effects of Missing Data Methods in SEM Under Conditions of Incomplete and Nonnormal Data. The Journal of Experimental Education 85, 231–258. ISSN: 0022-0973. 10.1080/00220973.2015.1134418 (2024) (Apr. 3, 2017).

18. Bassett, E., Broadbent, J., Gill, D., Burgess, S. & Mason, A. M. Inconsistency in UK Biobank Event Definitions From Different Data Sources and Its Impact on Bias and Generalizability: A Case Study of Venous Thromboembolism. American Journal of Epidemiology 193, 787–797. ISSN: 0002-9262. 10.1093/aje/kwad232 (2024) (May 7, 2024).

19. Lorente, S., Viladrich, C., Vives, J. & Losilla, J.-M. Tools to Assess the Measurement Properties of Quality of Life Instruments: A Meta-Review. BMJ Open 10, e036038. ISSN: 2044-6055, 2044-6055. PMID: 32788186. https://bmjopen.bmj.com/content/10/8/e036038 (2025) (Aug. 1, 2020).

20. Ruyssen-Witrand, A., Tubach, F. & Ravaud, P. Systematic Review Reveals Heterogeneity in Definition of a Clinically Relevant Difference in Pain. Journal of Clinical Epidemiology 64, 463–470. ISSN: 0895-4356. https://www.sciencedirect.com/science/article/pii/S0895435610002593 (2025) (May 1, 2011).

21. Zhang, F. & Finkelstein, J. Inconsistency in Race and Ethnic Classification in Pharmacogenetics Studies and Its Potential Clinical Implications. Pharmacogenomics and Personalized Medicine 12, 107–123. ISSN: null. PMID: 31308725. https://www.tandfonline.com/doi/abs/10.2147/PGPM.S207449 (2024) (July 2, 2019).

22. Brewster, R. C. et al. Race and Ethnicity Reporting and Representation in Pediatric Clinical Trials. Pediatrics 151, e2022058552. ISSN: 0031-4005. 10.1542/peds.2022-058552 (2024) (Mar. 14, 2023).

23. Minicuci, N. et al. Data Resource Profile: Cross-national and Cross-Study Sociodemographic and Health-Related Harmonized Domains from SAGE plus CHARLS, ELSA, HRS, LASI and SHARE (SAGE+ Wave 2). International Journal of Epidemiology 48, 14–14j. ISSN: 0300-5771. 10.1093/ije/dyy227 (2024) (Feb. 1, 2019).

24. Neidhart, M. et al. A protocol for data harmonization in large cohorts. Nature Mental Health 2, 1134–1137 (2024).

25. Siddique, J., de Chavez, P. J., Howe, G., Cruden, G. & Brown, C. H. Limitations in Using Multiple Imputation to Harmonize Individual Participant Data for Meta-Analysis. Prevention Science 19, 95–108. ISSN: 1573-6695. 10.1007/s11121-017-0760-x (2024) (Feb. 1, 2018).

26. Curran, P. J. & Hussong, A. M. Integrative Data Analysis: The Simultaneous Analysis of Multiple Data Sets. Psychological Methods 14, 81–100. ISSN: 1939-1463, 1082-989X. https://doi.apa.org/doi/10.1037/a0015914 (2024) (2009).

27. Hoffmann, M. S. et al. Harmonizing bifactor models of psychopathology between distinct assessment instruments: reliability, measurement invariance, and authenticity. International Journal of Methods in Psychiatric Research 32, e1959 (2023).

28. Hoffmann, M. S. et al. An evaluation of item harmonization strategies between assessment tools of psychopathology in children and adolescents. Assessment 31, 502–517 (2024).

29. Marquand, A. F. et al. Learning latent profiles via cognitive growth charting in psychosis: design and rationale for the PRECOGNITION project. Schizophrenia Bulletin Open, sgaf007 (2025).

30. An, U. et al. Deep Learning-Based Phenotype Imputation on Population-Scale Biobank Data Increases Genetic Discoveries. Nature Genetics 55, 2269–2276. ISSN: 1546-1718. https://www.nature.com/articles/s41588-023-01558-w (2024) (Dec. 2023).

31. Buuren, S. van & Groothuis-Oudshoorn, K. Mice: Multivariate Imputation by Chained Equations in R. Journal of Statistical Software 45, 1–67. ISSN: 1548-7660. 10.18637/jss.v045.i03 (2024) (Dec. 12, 2011).

32. Jolani, S. Hierarchical Imputation of Systematically and Sporadically Missing Data: An Approximate Bayesian Approach Using Chained Equations. Biometrical Journal 60, 333–351. ISSN: 1521-4036. https://onlinelibrary.wiley.com/doi/abs/10.1002/bimj.201600220 (2024) (2018).

33. Yoon, J., Jordon, J. & Schaar, M. Gain: Missing data imputation using generative adversarial nets in International conference on machine learning (2018), 5689–5698.

34. Jäger, S., Allhorn, A. & Bießmann, F. A benchmark for data imputation methods. Frontiers in big Data 4, 693674 (2021).

35. Caton, S., Malisetty, S. & Haas, C. Impact of Imputation Strategies on Fairness in Machine Learning. Journal of Artificial Intelligence Research 74, 1011–1035. ISSN: 1076-9757. https://www.jair.org/index.php/jair/article/view/13197 (2024) (June 27, 2022).

36. Geurts, P., Ernst, D. & Wehenkel, L. Extremely Randomized Trees. Machine Learning 63, 3–42. ISSN: 1573-0565. 10.1007/s10994-006-6226-1 (2024) (Apr. 1, 2006).

37. Haatveit, B. et al. Intra-and inter-individual cognitive variability in schizophrenia and bipolar spectrum disorder: an investigation across multiple cognitive domains. Schizophrenia 9, 89 (2023).

38. Ambrosen, K. S. et al. A machine-learning framework for robust and reliable prediction of short- and long-term treatment response in initially antipsychotic-naı”ve schizophrenia patients based on multimodal neuropsychiatric data. Translational psychiatry 10, 276 (2020).

39. Jessen, K. et al. Patterns of cortical structures and cognition in antipsychotic-naı”ve patients with first-episode schizophrenia: a partial least squares correlation analysis. Biological Psychiatry: Cognitive Neuroscience and Neuroimaging 4, 444–453 (2019).

40. Nielsen, M. Ø. et al. Differential effects of aripiprazole and amisulpride on negative and cognitive symptoms in patients with first-episode psychoses. Frontiers in Psychiatry 13, 834333 (2022).

41. Gottlieb, A. et al. Cohort-Specific Imputation of Gene Expression Improves Prediction of Warfarin Dose for African Americans. Genome Medicine 9, 98. ISSN: 1756-994X. 10.1186/s13073-017-0495-0 (2025) (Nov. 24, 2017).

42. Auwerx, C., Sadler, M. C., Reymond, A. & Kutalik, Z. From Pharmacogenetics to Pharmaco-Omics: Milestones and Future Directions. Human Genetics and Genomics Advances 3. issn: 2666-2477. https://www.cell.com/hgg-advances/abstract/S2666-2477(22)00016-1 (2025) (Apr. 14, 2022).

43. Schouten, R. M., Lugtig, P. & Vink, G. Generating Missing Values for Simulation Purposes: A Multivariate Amputation Procedure. Journal of Statistical Computation and Simulation 88, 2909–2930. ISSN: 0094-9655. 10.1080/00949655.2018.1491577 (2024) (Oct. 13, 2018).

44. Hofert, M., Jackson, J. & Hagenbuch, N. Bernoulli Amputation 2407.18572 [math, stat]. http://arxiv.org/abs/2407.18572 (2024). Pre-published.

45. Morvan, M. L. & Varoquaux, G. Imputation for prediction: beware of diminishing returns. arXiv preprint arXiv:2407.19804 (2024).

46. John-Mathews, J.-M., Cardon, D. & Balagué, C. From Reality to World. A Critical Perspective on AI Fairness. Journal of Business Ethics 178, 945–959. ISSN: 1573-0697. 10.1007/s10551-022-05055-8 (2025) (July 1, 2022).

47. Goh, W. W. B., Yong, C. H. & Wong, L. Are Batch Effects Still Relevant in the Age of Big Data? Trends in Biotechnology 40, 1029–1040. ISSN: 0167-7799, 1879-3096. PMID: 35282901. https://www.cell.com/trends/biotechnology/abstract/S0167-7799(22)00036-1 (2025) (Sept. 1, 2022).

48. Marquand, A. F., Rezek, I., Buitelaar, J. & Beckmann, C. F. Understanding heterogeneity in clinical cohorts using normative models: beyond case-control studies. Biological psychiatry 80, 552–561 (2016).

49. Marquand, A. F. et al. Conceptualizing mental disorders as deviations from normative functioning. Molecular psychiatry 24, 1415–1424 (2019).

50. De Boer, A. A. et al. Non-Gaussian normative modelling with hierarchical Bayesian regression. Imaging Neuroscience 2, 1–36 (2024).

51. Breiman, L. Random Forests. Machine Learning 45, 5–32. ISSN: 1573-0565. 10.1023/A:1010933404324 (2024) (Oct. 1, 2001).

52. Shah, A. D., Bartlett, J. W., Carpenter, J., Nicholas, O. & Hemingway, H. Comparison of Random Forest and Parametric Imputation Models for Imputing Missing Data Using MICE: A CALIBER Study. American Journal of Epidemiology 179, 764–774. ISSN: 0002-9262. 10.1093/aje/kwt312 (2024) (Mar. 15, 2014).

53. Shadbahr, T. et al. The Impact of Imputation Quality on Machine Learning Classifiers for Datasets with Missing Values. Communications Medicine 3, 1–15. ISSN: 2730-664X. https://www.nature.com/articles/s43856-023-00356-z (2024) (Oct. 6, 2023).

54. Simonsen, C. et al. Neurocognitive dysfunction in bipolar and schizophrenia spectrum disorders depends on history of psychosis rather than diagnostic group. Schizophrenia bulletin 37, 73–83 (2011).

